# Assessing the Population Consequences of Disturbance and Climate Change for the Pacific Walrus

**DOI:** 10.1101/2023.10.12.562073

**Authors:** Devin L. Johnson, Joseph M. Eisaguirre, Rebecca L. Taylor, Joel L. Garlich-Miller

## Abstract

Climate change and anthropogenic disturbance are increasingly affecting wildlife at a global scale. Predicting how varying types and degrees of disturbance may interact to influence population dynamics is a key management challenge. Population Consequences of Disturbance (PCoD) models provide a framework to link effects of anthropogenic disturbance on an individual’s behavior and physiology to population-level changes. Bioenergetic models often constitute the basis of these frameworks, wherein an individual’s daily energy balance is simulated over the course of its lifetime, allowing many individuals to be subjected to different environmental conditions and ultimately simulate population-level vital rates under varying degrees of disturbance. In the present study, we develop a Pacific walrus (*Odobenus rosmarus divergens*) PCoD model to encompass the population-level effects of both anthropogenic disturbance and climate change. Pacific walruses are an Arctic/subarctic ice-associated pinniped. As the Arctic has become increasingly ice-free with climate change, walruses spend more time on land-based (rather than ice-based) haulouts from which they must expend more energy to reach foraging areas, and where they have a greater risk of predation and disturbance-based mortalities. Concurrently, sea ice loss is increasing the anthropogenic footprint in Arctic regions (e.g., fisheries, shipping, energy exploration) which creates additional disturbance. We developed a bioenergetic Dynamic Energy Budget (DEB) model for the Pacific walrus and applied it to four scenarios (ranging from optimistic-pessimistic) which incorporate different global sea ice model projections along with varying degrees of anthropogenic disturbance. All scenarios indicated a decline in Pacific walrus carrying capacity and population growth rate (and thus overall abundance) to the end of the 21^st^ century, but demonstrated that the intensity of that decline could be mitigated by global efforts to reduce carbon emissions (i.e., lessening the rate of sea ice loss) and local management and conservation efforts to protect sensitive habitat areas. In summary, we introduced a flexible PCoD modelling framework in a novel context which will prove useful to researchers studying walruses and other species similarly threatened by rapid environmental change.

## INTRODUCTION

Under changing global conditions, wildlife populations are under increasing pressure from direct and indirect anthropogenic effects that influence animal behavior on an individual basis. These behavioral changes can alter an animal’s physiology and lifetime reproductive success, often with lasting population-level effects (e.g., Pirotta et al., 2019). Thus, frameworks are needed for analyzing and predicting how wild animals respond to disturbance and changing environmental conditions, and how those responses result in population changes for species of conservation and management concern.

Population Consequence of Disturbance (PCoD) models (reviewed by Pirotta et al., 2018) provide a framework to link disturbance at the individual level to effects at the population level. Since its inception (Middleton et al., 2013), this approach has been applied to a wide range of animal populations and has recently been adapted to include interactive effects of both disturbance and environmental change (Pirotta et al., 2019). Bioenergetic models are particularly useful in this framework because they can simulate individual movement and energy acquisition—and subsequent effects on body condition, survival, and reproduction (e.g., New et al., 2013). The Dynamic Energy Budget (DEB; De Roos et al., 2009) is one such model designed to simulate a female marine mammal’s daily body condition and lifetime reproductive success by balancing energy gain (resource assimilation) and energy loss (metabolism, growth, fetal development, and lactation). Hin et al. (2019) further integrated a DEB for use in a PCoD framework for pilot whales, introducing a powerful tool for simulating population consequences of disturbance and environmental change in marine mammal populations.

The Pacific walrus (*Odobenus rosmarus divergens*) is a large, tusked pinniped distributed across the Bering and Chukchi seas and coasts of Alaska and Russia. They are estimated to have a population of 160,000–354,000 individuals (95% CrI; Beatty et al., 2022) and have been an important traditional subsistence resource for local indigenous communities for millennia (MacCracken, 2012). As an Arctic/subarctic specialist that depends on sea ice for its reproductive success (Fay, 1982), the Pacific walrus population may decline to the end of the 21^st^ century (e.g., MacCracken et al., 2017). As the climate has warmed and sea ice has become less available for female Pacific walruses and their calves to rest on in the Chukchi sea in summer and autumn, they have increasingly hauled out to rest on land (Fischbach et al. 2022), where they run a greater risk of disturbance-based mortality (Jay et al., 2012, Udevitz et al., 2013). Additionally, they spend less time foraging and resting when sea ice is not available, because of the long distance between some land-based haulouts and productive foraging areas (Jay et al. 2017). As the region becomes increasingly ice-free, human presence, development, and impacts are also likely to increase (e.g., Lavelle, 2013; Melia et al., 2016; Huntington et al., 2020). Indigenous Knowledge (IK) holders (i.e., members of subsistence walrus hunting communities) have raised concerns about impacts of increasing direct anthropogenic disturbance such as oil and gas activities, ship and air traffic, and commercial fisheries on the Pacific walrus population (MacCracken et al., 2017; Table S1). Although several of these factors have been addressed individually (e.g., Udevitz et al., 2013; Udevitz et al., 2017), there is need for a comprehensive framework which can assess the overall impact of environmental change and disturbance.

The primary objective of this study was to develop a framework for simulating the consequences of climate change and anthropogenic disturbance on the Pacific walrus population. We developed a DEB (following Hin et al., 2019) applying walrus-specific physiological parameters when available, and relying on parameter values and assumptions from other marine mammals where walrus-specific data were unavailable. By incorporating sea ice projections in a PCoD framework, we predicted population-level responses to an array of climate change and disturbance scenarios. This tool should prove useful for evaluating conservation and management options for a species in a complex, dynamic system.

## METHODS

### OVERVIEW

We developed a Dynamic Energy Budget (DEB) for the Pacific walrus, based on a set of bioenergetic parameters that simulates a female walrus’ energy balance and reproductive success over the course of her lifetime. The DEB incorporates estimates of walrus seasonal movement and activity budgets (and associated energy expenditure) in response to sea ice cover. The baseline model (i.e., under contemporary sea ice conditions) was calibrated to match population-level estimates of reproductive and age-specific survival rates from the most recent available data (Taylor et al., 2018). We developed a suite of combined climate/disturbance scenarios that incorporate sea ice projections from recent global climate models, varying degrees of anthropogenic disturbance and mitigation of mortality at coastal haulouts, and potential changes to prey density. After we calibrated the baseline DEB, we applied each scenario in a PCoD framework to predict the response of the Pacific walrus population to different potential conditions up to the middle and end of the 21^st^ century.

### DYNAMIC ENERGY BUDGET

The DEB (e.g., De Roos et al., 2009; Hin et al. 2019) is a state-specific bioenergetic model that tracks the daily energy assimilation and expenditure of female marine mammals to ultimately estimate lifetime reproductive success. We developed a DEB model that relies on parameter values from the Pacific walrus literature, or when these were unavailable, from the best available surrogate species (Table 1).

**Table 1:** Summary of model parameter values and their sources. Values are walrus-specific unless otherwise noted.

| Symbol | Units | Value | Definition | Source |
| --- | --- | --- | --- | --- |
| **Age Parameters** |  |  |  |  |
| $T_{MAX}$ | days | 16060 | Maximum walrus age (44 years) | Taylor et al. (2015) |
| $T_{RMIN}$ | days | 1460 | Minimum possible reproductive age (4 years) | Fay (1982) |
| $T_{RMAX}$ | days | 10949 | Maximum reproductive age (30 years) | Fay (1982) |
| $T_P$ | days | 310 | Duration of pregnancy (active gestation) | Noren et al. (2014) |
| $TC_R$ | days | 450 | Calf age at which female begins to reduce milk supply and the calf's foraging efficiency is 50% of an adult's foraging efficiency | Derived from data in Fay (1982) |
| $TC_W$ | days | 730 | Calf age at weaning (2 years) | Noren et al. (2014) |
| **Growth Parameters** |  |  |  |  |
| $L_0$ | cm | 113 | Length at birth | Fay (1982) |
| $L_\infty$ | cm | 280 | Maximum length | Fay (1982) |
| $\nu$ | $\text{cm}\cdot\text{day}^{-1}$ | $6.85\cdot 10^{-4}$ | Von Bertalanffy growth rate | Garlich-Miller & Stewart (1998)<sup>‡</sup> |
| $\omega_s$ | $\text{kg}\cdot\text{cm}^{-1}$ | $1.82\cdot 10^{-4}$ | Structural mass-length scaling constant | Garlich-Miller & Stewart (1998)<sup>‡</sup> |
| $\omega_e$ | — | 2.72 | Structural mass-length scaling exponent | Garlich-Miller & Stewart (1998)<sup>‡</sup> |
| **Metabolic Parameters** |  |  |  |  |
| $\sigma_G$ | $\text{MJ}\cdot\text{kg}^{-1}$ | 28.5 | Energy cost per unit of growth in structural mass | Noren et al. (2014) |
| $k$ | $\text{MJ}\cdot\text{kg}^{-1}\cdot\text{day}^{-1}$ | $70\cdot\text{MASS}^{0.75}_*$ | Mass-Specific Basal Metabolic Rate | Kleiber (1975) |
| $\sigma_{M\_LI}$ | $\text{L O}_2\cdot\text{min}^{-1}$ | $0.00101\cdot\text{MASS}^{1.25}_*$ | Mass-Specific Field Metabolic Rate for resting on Land or Ice | Rode et al. (2023) |
| $\sigma_{M\_WS}$ | $\text{L O}_2\cdot\text{min}^{-1}$ | $0.00245\cdot\text{MASS}^{1.13}_*$ | Mass-Specific Field Metabolic Rate for activity in Water - Swimming | Borque-Espinosa et al. (2021) |
| $\sigma_{M\_WD}$ | $\text{L O}_2\cdot\text{min}^{-1}$ | $0.02820\cdot\text{MASS}^{0.73}_*$ | Mass-Specific Field Metabolic Rate for activity in Water - Diving | Borque-Espinosa et al. (2021) |
| $\sigma_{M\_WR}$ | $\text{L O}_2\cdot\text{min}^{-1}$ | $0.00123\cdot\text{MASS}^{1.22}_*$ | Mass-Specific Field Metabolic Rate for activity in Water - Resting | Borque-Espinosa et al. (2021) |
| **Resource Parameters** |  |  |  |  |
| $R$ | $\text{MJ}\cdot\text{m}^3$ | 4.034 | Resource availability | This study |
| $\varepsilon_+$ | $\text{MJ}\cdot\text{kg}^{-1}$ | 32 | Anabolic reserve conversion efficiency | Udevitz et al. (2017) |
| $\varepsilon_-$ | $\text{MJ}\cdot\text{kg}^{-1}$ | 32 | Catabolic reserve conversion efficiency | Udevitz et al. (2017) |
| $\rho_s$ | - | 0.10 | Starvation threshold (reserve mass/total body mass) | Noren et al. (2009)** |
| $\mu_s$ | - | $0.2^{\dagger}$ | Starvation mortality scalar | Hin et al. (2019) |
| $\gamma$ | — | $3^{\dagger}$ | Shape parameter for relationship between resource assimilation and age | Hin et al. (2019) |
| $\rho$ | — | 0.3 or 0.39 | Target body condition for normal and late pregnancy (reserve mass/total body mass). | Harwood et al. 2019 |
| $\eta$ | — | $15^{\dagger}$ | Slope of assimilation response around target body condition | Hin et al. (2019) |
| **Lactation Parameters** |  |  |  |  |
| $\sigma_L$ | — | $0.86^{\dagger}$ | Lactation efficiency | Lockyer (1993) |
| $\Phi_M$ | - | 4.034 | Milk energetic scaling parameter | This study |
| $\xi_M$ | — | $-2^{\dagger}$ | Shape parameter for relationship between milk provisioning and female body condition | Hin et al. (2019) |
| $\xi_C$ | — | $0.25^{\dagger}$ | Shape parameter for relationship between milk assimilation and calf age | Hin et al. (2019) |

\* MASS refers to the individual's total mass, defined as $S_t$ (structural mass) + $F_t$ (reserve mass).
\*\*Value for Steller sea lions (*Eumatopias jubatus*)
<sup>†</sup>Value assumed in a DEB model developed for North Atlantic long-finned pilot whales
<sup>‡</sup>Value from a study on Atlantic walrus (*Odobenus Rosmarus*) rather than the Pacific walrus subspecies

Individual females are simulated beginning from weaning age (*TC_W_* = 2 years) until they die, either from old age (*T_MAX_* = 44 years), or from other causes of mortality (see *Survival*). A female can occupy one of four states at any given time: “non-reproductive” (either an adult who is neither pregnant nor lactating or a juvenile from weaning to age of first implantation), pregnant (but not lactating), lactating (but not pregnant), and lactating and pregnant. Each state confers different energy requirements depending upon the body mass of the individual and the growth stage of her fetus or calf (if applicable). Our series of interrelated state-dependent equations that balance daily energy input and expenditures within the DEB is conceptualized in Figure 1.

**Figure 1.**
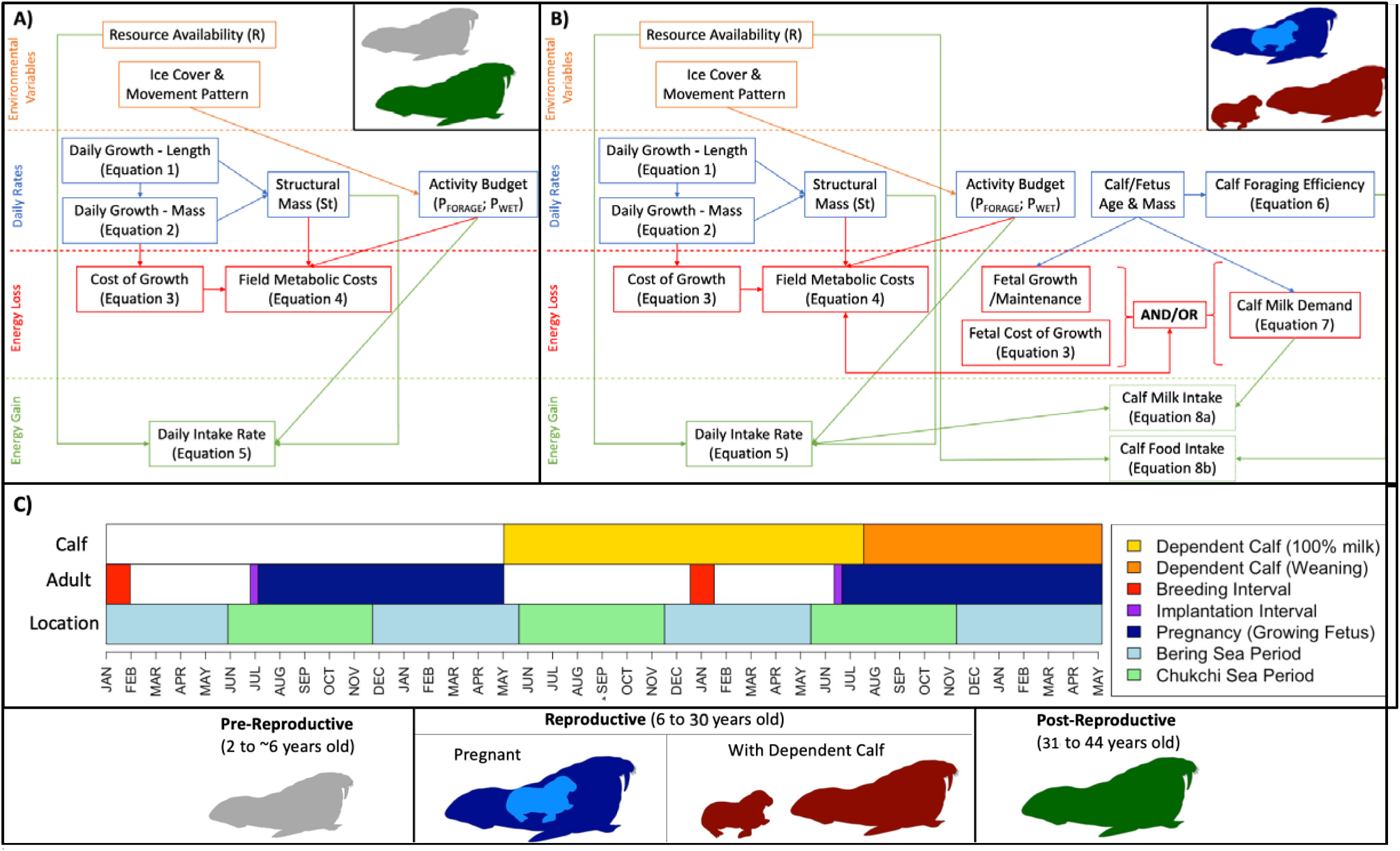
Conceptual diagram of the DEB model structure for A) non-reproductive and B) reproductive adult female Pacific walruses, denoting the factors contributing to daily energy balance. Some nodes in panel B depend on whether the female is pregnant, has a dependent calf, or both. A reproductive-age female who fails to implant or has a fetal or calf mortality would also fall under state A. Panel C) depicts the phenology of a complete reproductive cycle of a female walrus as considered in the model (i.e., from breeding to calf independence), assuming she is able to successfully become pregnant and raise young in an optimum fashion. More detail on model parameters shown here can be found in Table 1.

### Structural Mass

In this section we characterize Pacific walrus “structural mass” (*S_t_*; i.e., tissue such as bones & organs that cannot be catabolized for energetic needs), whereas “reserve mass” (*F_t_*; i.e., muscle and blubber) is modified in the sections below pertaining to growth, metabolism, and reproduction. Because we require structural mass (as opposed to total mass) for the DEB model, we were unable to directly apply documented mass growth curves (Fay, 1982) and instead relied on the generalized von Bertalanffy growth equation, which provided a reasonable fit to historical, empirical data for Pacific walruses (McLaren, 1993):

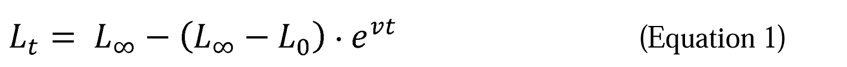

In Equation 1: *L_t_* is body length at age *t* (in days); *L*_∞_ is the maximum possible length (280 cm); *L_0_*is the length at birth (113 cm); and *v* is the growth rate (0.000685 cm·day^-1^; Table 1). Body length was then incorporated into the following equation to estimate structural mass at age *t* (*S_t_*):

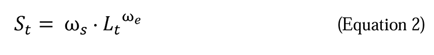

In Equation 2: ω_s_ is the structural mass-length scaling constant (1.82·10^-4^ kg·cm^-1^, Table 1); and ω_e_is the structural mass-length scaling exponent (2.72; Table 1). We assumed male and female calves grow at the same rate from *T_0_* (birth day) to *TC_W_* (weaning day). Although walruses are sexually dimorphic, sex-specific differences in body length growth curves appear minimal, and mass-length relationships are not significantly different between sexes in the first two years of life (Fay 1982; Garlich-Miller & Stewart, 1998).

We assumed fetuses grow at a constant rate from a length of 0 cm (at implantation) to *L_0_* (at birth), over the course of active gestation (*T_P_* = 310 days; Table 1; Fay 1982). We estimated the structural mass of the fetus using the same mass-length relationship that was used for females. The resulting age-specific growth curves for female and fetal walruses are displayed in Figure S1.

### Energy Intake & Assimilation

Each female walrus assimilates energy each day from feeding according to the following equation:

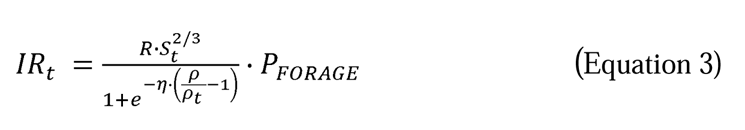

In Equation 3: *IR_t_* is the intake rate on day *t* (MJ/day); *R* is the resource availability parameter; *P_FORAGE_* (the proportion of the day the walrus is foraging) is drawn from a probability distribution that varies by region and ice cover (see *<u>POPULATION CONSEQUENCES OF</u> <u>CLIMATE CHANGE</u>*); *S_t_*is the animal’s structural mass; η is the slope of the assimilation response around the target body condition (15, Table 1); ρ is the target body condition (0.3 or 0.39; Table 1), and ρ*_t_* is the animal’s relative body condition (reserve mass/total mass, *F_t_*/*W_t_*). We define resource availability (*R*) as the daily amount of metabolizable energy each female has available to them from food resources. Thus, *R* is a generalized metric of environmental quality that is not intended to directly reflect empirical estimates of benthic biomass (e.g., Wilt et al., 2014), but instead allows for scaling with ice-associated activity budgets (the proportion of the day each walrus spends foraging) and model calibration. Equation 3 assumes individuals forage at the maximum possible efficiency when their relative body condition (ρ*_t_*) is low (i.e., near their starvation threshold, ρ*_s_*, Table 1), and reduce their foraging efficiency when they approach their target body condition (ρ), based on a slope parameter η (following Hin et al., 2019). The target body condition parameter (ρ) is set to 0.3 during most circumstances but rises to 0.39 during the second half of pregnancy to replicate an observed increase in the reserves of pregnant females (Noren et al., 2014). Following this equation, animals are allowed to compensate for the effect of lost foraging opportunities on their relative body condition by increasing energy assimilation on subsequent days, provided sufficient resources are available (Fig S2).

In many marine mammals, energy reserves (e.g., blubber & muscle) provide a buffer for incoming and outgoing energy flows for both the female and her calf (De Roos et al., 2009). In the model, surplus energy is converted to reserves if the assimilated energy (*IR_t_*) exceeds total energy expenditure on a given day; reserves are catabolized if the opposite is true. The energetic cost of creating each kg of reserve tissue is ε*^+^* MJ, and each kg of tissue provides ε*^-^* MJ when catabolized (Table 1).

Walrus calves have foraging skills that are poor at birth, increase with age, and asymptote later in life (Fay, 1982). We modelled this (following the relationship assumed by Hin et al. (2019)) as:

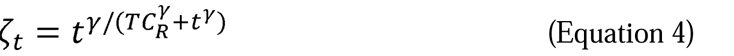

In Equation 4: is foraging efficiency on day *t*; γ is a shape parameter quantifying the relationship between intake rate and age for calves 0–3 years of age (3; Table 1); and *TC_R_* is the age at which calf foraging efficiency is 50% of adult foraging efficiency (450 days; Table 1).

Calves rely wholly or partially on milk until they are weaned (*TC_W_* = 730 days of age) following the relationship:

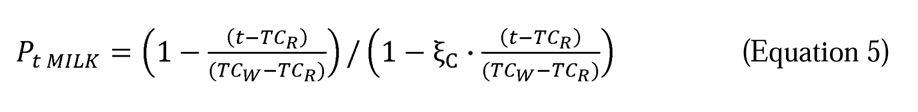

In Equation 5: *P_t MILK_* is the proportion of a calf’s energy requirement that comes from milk on day *t*; *T_CR_*is the calf age at which its mother begins to reduce milk supply (450 days); *T_CW_* is the age at which a calf is fully weaned (730 days); and ξ_C_ is a shape parameter for the relationship between milk assimilation and calf age based on pilot whales (0.25; Table 1).

The calf intake rate (the total amount of energy a calf obtains, Fig S3) is derived from the following equation in a similar fashion to Equation 3, with separate terms for assimilation from milk and foraging:

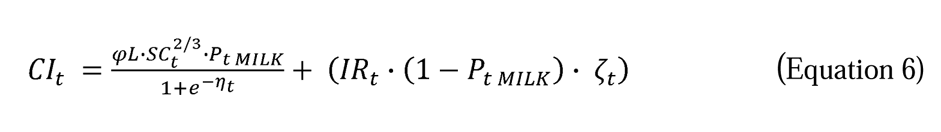

In Equation 6: *CI_t_*is the calf’s milk intake rate on day *t*, φ*_L_* is a scaler that reflects the energy density of milk (i.e., the equivalent to *R* from Eqn. 3); *SC_t_* is the calf’s body mass at its age on day *t* (Equation 2); *IR_t_* is the mother’s resource intake (Equation 3); and, is the calf’s foraging efficiency at its age on day *t* (Equation 4).

If the mother’s body condition reaches or approaches its starvation threshold (ρ*_s_*), she reduces the amount of milk she supplies to her calf (ψ*_t_*, Fig S3) following the equation:

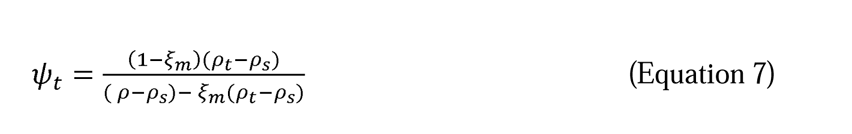

In Equation 7: ξ*_m_* is a shape parameter (-2; Table 1); and ρ*_t_,* ρ, and ρ*_s_* represent the mother’s body condition (current, targeted, and starvation threshold, respectively).

With these values, if the mother is in good body condition (i.e., ρ*_t_* ≃ 0.3), she provides all of the calf’s demands for milk during the first 14 months of its life, then milk provisioning falls monotonically to zero at weaning as the calf begins to forage for a larger percentage of its energy demand. If females are in very poor condition, they will cease to provide any milk for their calves before they are weaned, often resulting in calf mortality.

The total daily energetic cost of a female’s milk production can be calculated by dividing her calf’s energy intake from milk on day *t* (the first term of Equation 6) by the efficiency at which reserves are converted to milk (σ*_L_*= 0.86; Table 1). These costs are highest during the first year of lactation (when the entire calf energy requirement comes from milk) and particularly high during the first 50 days of lactation as the calf grows most rapidly (Fig. S3). Because simulated females under 10 years of age have lower reserves than older females, they are more likely to have starvation-related calf mortalities, mimicking relationships observed in wild populations (Noren et al., 2014).

### Cost of Growth & Metabolism

The daily cost of growth is calculated as the difference in structural mass between consecutive days, multiplied by the energy cost per unit of structural growth (σ*_G_*= 28.5 MJ·kg^-1^; Table 1):

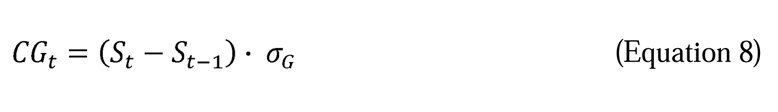

Along with growth, each animal’s daily field metabolic cost (*FM_t_*) is calculated based upon its daily activity budget. A simulated walrus can occupy each of four activity states which confer different mass-specific metabolic rates (Table 1): resting on land or ice (σ_M__LI); surface/subsurface swimming (σ_M__WS); diving (i.e., foraging on the ocean floor; σ_M__WD); or resting in the water (σ_M__WR; Fig. S4). Each animal spends a certain percentage of each day in the water (*P_WATER_*), and a certain percentage of its in-water time actively foraging (*P_FORAGE_*). When foraging, a walrus spends a certain amount of its time either diving (77.2%) or resting between dives (22.8%; mean values from Udevitz et al., 2017). We assume when walruses are in the water and not foraging, they are either traveling to foraging grounds or migrating. From this we can calculate an animal’s overall metabolic rate of being in the water (σ*_M__W*), and multiply those metabolic rates by structural mass (following Kleiber 1975) to calculate total daily field metabolic costs (*FM_t_*):

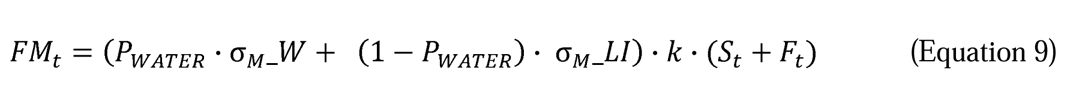

where

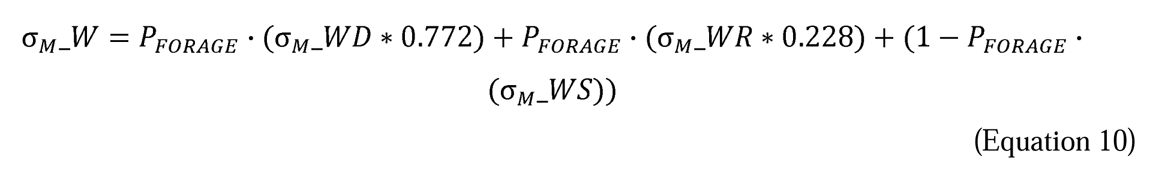

In Equations 9 and 10: *k* is the mass-specific basal metabolic rate (Table 1) that is multiplied by the activity scalers (<u>σ</u>_M_, Table 1); *S_t_* is the animal’s structural mass on day *t*; and *F_t_* is the animal’s reserve mass. The daily activity budget of neonatal calves (age 0–90 days) is different from that of all other walruses because they are thought to spend the majority of their time resting as they rapidly accumulate mass from their mother’s milk (Fay 1982). Thus, the model applies resting metabolic rates to calves during this period. Calves > 90 days of age were assumed to have the same activity budgets as their mothers, but the mass-specific metabolic scalers modified the energetic costs of this activity budget based on the calf’s size.

### Reproduction

If a simulated female is over four years of age (*T_RMIN_*; Table 1) and her body condition exceeds a certain threshold (ρ > 0.3; Harwood et al. 2019) during a 10-day period starting on July 1 (mean implantation date; Fay, 1982), she is capable of becoming pregnant. We applied age-specific ovulation rates to each female from years 4–9 based on published estimates (Fay 1982), ranging from 10.7% at age 4 to 100% at age 10. We incorporate these age-specific ovulation rates multiplied by a 90% implantation success rate to determine the probability a female becomes pregnant (following Fay, 1982). Pacific walruses are only physiologically capable of producing one calf every two years due to a gestation period (passive + active) that is longer than a year (Noren et al., 2014), thus the model assumes females cannot become pregnant during their first year of lactation. Females were given a maximum reproductive age (*T_RMAX_*) of 30 years (following Fay et al., 1982), allowing for a 24-year potential reproductive interval. To summarize results throughout this study we define a “reproductive adult” to be between 6 and 30 years of age, inclusive, although the DEB includes the small probability that a walrus is capable of becoming pregnant at ages 4 or 5 (following Fay, 1982).

During pregnancy, a female must cover the costs of fetal growth and maintenance (Equation 8) for the fetus to survive (reviewed in McHuron et al., 2023). Thus, the mass of the growing fetus is added to the structural mass of the pregnant female, and fetal maintenance is included in her overall metabolic cost (*FM_t_*), applying a resting metabolic rate to the fetus’s body mass. Additional caloric requirements for pregnant females are accounted for by the target body condition parameter (ρ) which rises from 0.3 to 0.39 during the second half of pregnancy (see *Energy Intake and Assimilation*). Provided the pregnant female’s body condition stays above the starvation threshold (ρ*_s_*), we assume she transfers sufficient energy to her fetus for it to grow. Additionally, fetal survival was assumed to be 95% (in the absence of starvation events; following Fay 1982), thus we applied a daily fetal survival rate of^334^√0.95 = 0.99985 over the course of the pregnancy.

### Survival

We derived an age-dependent cumulate survival curve (for survival from background mortality causes, Fig. S5) by building a piecewise function based on age-specific survival estimates from a Pacific walrus integrated demographic population model (IPM; the most parsimonious model from Taylor et al. 2018). This curve employs five age classes: neonates (age 0–90 days, annual survival = 0.65); older calves (age 91–365 days, annual survival = 0.76); juveniles (age 1–5 years, annual survival = 0.90); reproductive adults (age 6–30 years; annual survival = 0.99); and post-reproductive adults (age 31–44 years, annual survival = 0.55).

The age at death under baseline (good) conditions for each individual (female or calf) was set by drawing a random number between 0 and 1, and death occured when the cumulative survival probability (Fig. S5) fell below this value. Any simulated individual alive according to this baseline age at death could still die due to poor body condition. Thus, the mortality rate for each simulated individual increased when its relative body condition fell below a pre-defined starvation threshold (ρ*_s_*). On each such day when ρ*_t_* < ρ*_s_*, a Bernoulli trial was undertaken to determine the individual’s survival with probability,*_t_* defined by the following equation:

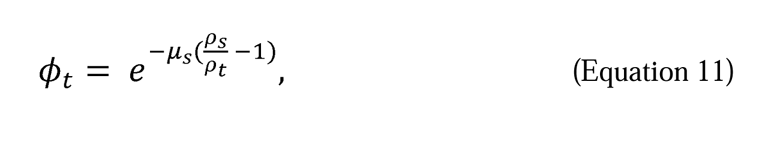

where μ_s_ is a scaler defining the strength of the starvation mortality relationship (0.2; Table 1).

### Population Growth Rate and Intrinsic Rate of Increase

Individuals simulated in the DEB represent a random sample of all possible female life histories, and their mean reproductive success and mean age of death (life expectancy) can be used to derive an estimate of population growth rate (Harwood et al., 2019).

By simulating many (e.g., 10,000 individuals) we were able to estimate population-level parameters under different environmental conditions and scenarios (Hin et al., 2019). We defined population-level lifetime reproductive success (hereafter “reproductive success”: *S_r_*) as the mean number of female offspring (assuming a 50:50 calf sex ratio) that were successfully raised to weaning age by each simulated female during her lifetime. The annual population growth rate (λ) can then be estimated using the following equation (e.g., Turchin, 2003):

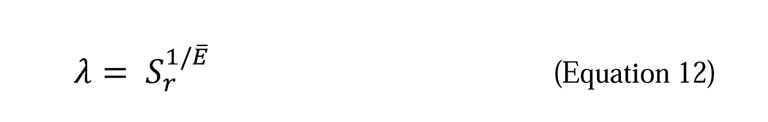

In Equation 12, e^-^ is life expectancy in years, and *S_r_* is the mean reproductive success of females in the population, assuming a random sample of female life histories. Given λ, we can estimate the intrinsic rate of increase (*r* = ln(λ)). Finally, the maximum intrinsic rate of increase (*r_max_*) is simply the rate of population increase when the population is unlimited by resources. Thus, we can estimate *r_max_* by effectively providing unlimited resources to the simulation (e.g., multiplying *R* [resource availability; Equation 3] by 10) and then calculating *r* (Harwood et al., 2019). We compared this method against other conventional equations for deriving *r_max_* (Cortes, 2018), and found it fell within the range of those estimates (Fig. S6) and produced values we would expect based on the species’ life history (e.g., Moore et al., 2018; Romero et al., 2017).

### Model Calibration

One benefit of the DEB modeling framework is that it involves flexible resource availability parameters (*R* and φ*_L_*), which can be adjusted to simulate different environmental conditions. These are the primary mechanisms through which a baseline model can be calibrated such that its output matches empirical estimates (e.g., *N/K*, reproductive and age-specific survival rates). Once a baseline model is produced that matches estimated population parameters, it is possible to introduce disturbance and climate change scenarios in a PCoD framework to estimate predicted changes to those population parameters. We calibrated our model using the estimates of *r* and *r_max_*, along with an estimated population size (*N*) to derive carrying capacity (*K*) using the following formula (e.g., Turchin, 2003):

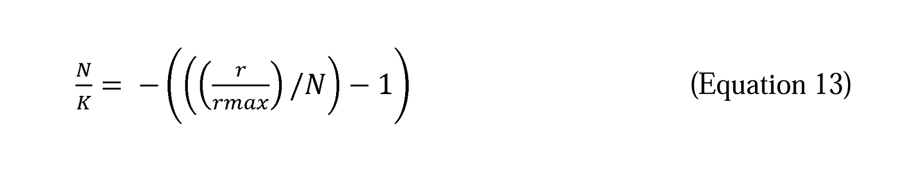

We calibrated our default DEB model with the values found in Table S2. We defined carrying capacity (*K*) as the point at which population size remains constant (i.e., *N_t_* _ *N_t_*_+1_). Pacific walrus population size (*N*; adult females) was most recently estimated in a genetic mark-recapture study (Beatty et al., 2022), at which time the population was considered near carrying capacity based on an IPM that indicated the population size was constant (Taylor et al. 2018, most parsimonious model). However, this conclusion was based on extant conditions that included harvest mortality; thus, some level of population growth would be expected if harvest was reduced (MacCracken et al., 2017). In this context, we calibrated the model to an *N*/*K* value of 0.9 to account for the effect of harvest on population dynamics, and *K* represents the size at which the population would remain constant over time if there were no harvest. We also calibrated the DEB to match density dependent annual reproductive rates (the annual probability a reproductive adult female gave birth to a female calf) and calf survival rates from the IPM (Taylor et al. 2018, most parsimonious model) (Fig. S7; Table S2).

### POPULATION CONSEQUENCES OF CLIMATE CHANGE

The primary environmental driver for climate-associated impacts on the Pacific walrus population is a reduction in sea ice availability during the summer/autumn months in the Chukchi sea. This has already influenced walrus movement and behavior in late summer and early autumn: in all but 2 years since 2007, lack of sea ice has forced females and young that traditionally haul out on sea ice near productive foraging areas to haul out by the tens of thousands on the northwest coast of Alaska (Fischbach et al. 2022). These behavioral changes have already influenced walrus activity budgets and thus energy expenditure (Jay et al., 2017), and may also influence demographic rates (MacCracken, 2012). Udevitz et al. (2017) developed a model that relates changes in sea ice availability to adult female walrus movements and activity budgets in the Chukchi Sea in summer/autumn, and then uses these to predict seasonal changes in body condition under different climate scenarios. In the present study, we extend this framework to also include effects of sea ice availability in the Bering Sea in winter/spring, and we use these year-round effects to predict population-level effects of climate change-induced sea ice loss.

Female and juvenile Pacific walruses traditionally migrate north to the Chukchi Sea in May and June, following the receding ice edge throughout the summer months before returning (with the sea ice) to the Bering Sea in October and November (Fay, 1982). Adult males typically stay in the Bering Sea year-round, using terrestrial haulouts during summer months (Garlich-Miller & Jay, 2000). Summer/autumn sea ice availability in the Chukchi Sea is already declining rapidly and is predicted to lead to months-long ice-free periods within the next 20 years—even under optimistic climate change scenarios (Wang & Overland, 2015). Previously, winter sea ice in the Bering Sea was expected to persist to the end of the 21^st^ century, even under pessimistic climate change conditions (Udevitz et al., 2017). However, the Intergovernmental Panel on Climate Change’s sixth assessment report features an updated set of global climate models (CMIP6; Coupled Model Intercomparison Project), which includes sea ice projections under two shared socio-economic pathways (ssp245 & ssp585; Fox-Kemper et al., 2021). These two pathways are commonly used by policymakers to characterize climate change: ssp245 (analogous to the previous RCP 4.5) represents an intermediate global carbon emissions scenario; whereas ssp585 (analogous to previous RCP 8.5) represents a pessimistic (“business as usual”) scenario where global carbon emissions continue to rise (O’Neill et al., 2016). These updated sea ice models predict greater losses to Bering Sea ice, projecting that it will be nearly ice-free year-round by 2100 according to a pessimistic (ssp585) trajectory (Fig. S8).

Udevitz et al. (2017) used walrus telemetry data from June–November 2008–2014 and contemporaneous ice cover estimates (from the National Ice Center daily Marginal Ice Zone [MIZ] and weekly Sea Ice Grid 3 [SIGRID-3] charts [https://usicecenter.gov/Products]) to describe five walrus movement patterns [A–E] among four regions of the Chukchi Sea [1–4]; Fig. 2). Each movement pattern conferred a different daily probability of transitioning between regions, and each region conferred a different daily probability of activity types (i.e., percentage of time spent in the water, and percentage of time spent foraging if in the water) as a function of the amount of sea ice cover within the current region (Udevitz et al. 2017). Within this model, these ice-related activity budgets were ultimately tied to Pacific walrus energy requirements and acquisition. In model projections, Udevitz et al. (2017) randomly assigned each simulated walrus in each summer to one movement pattern and one sequence of daily ice conditions obtained from one of the seven years of observed data (for current conditions), or from one year within each projected time frame from one climate projection model (for projected future conditions). We followed protocols outlined in Udevitz et al. (2017) for sea ice and sea ice-based movement projections, with three exceptions. First, we projected sea ice cover using updated CMIP6 models. Second, we included the Bering Sea from December–May (as well as the Chukchi Sea from June–November) to encompass the entire annual cycle of female Pacific walruses. Third, we tracked each individual simulated walrus for multiple years, over which we employed a total of six possible movement scenarios.

**Figure 2.**
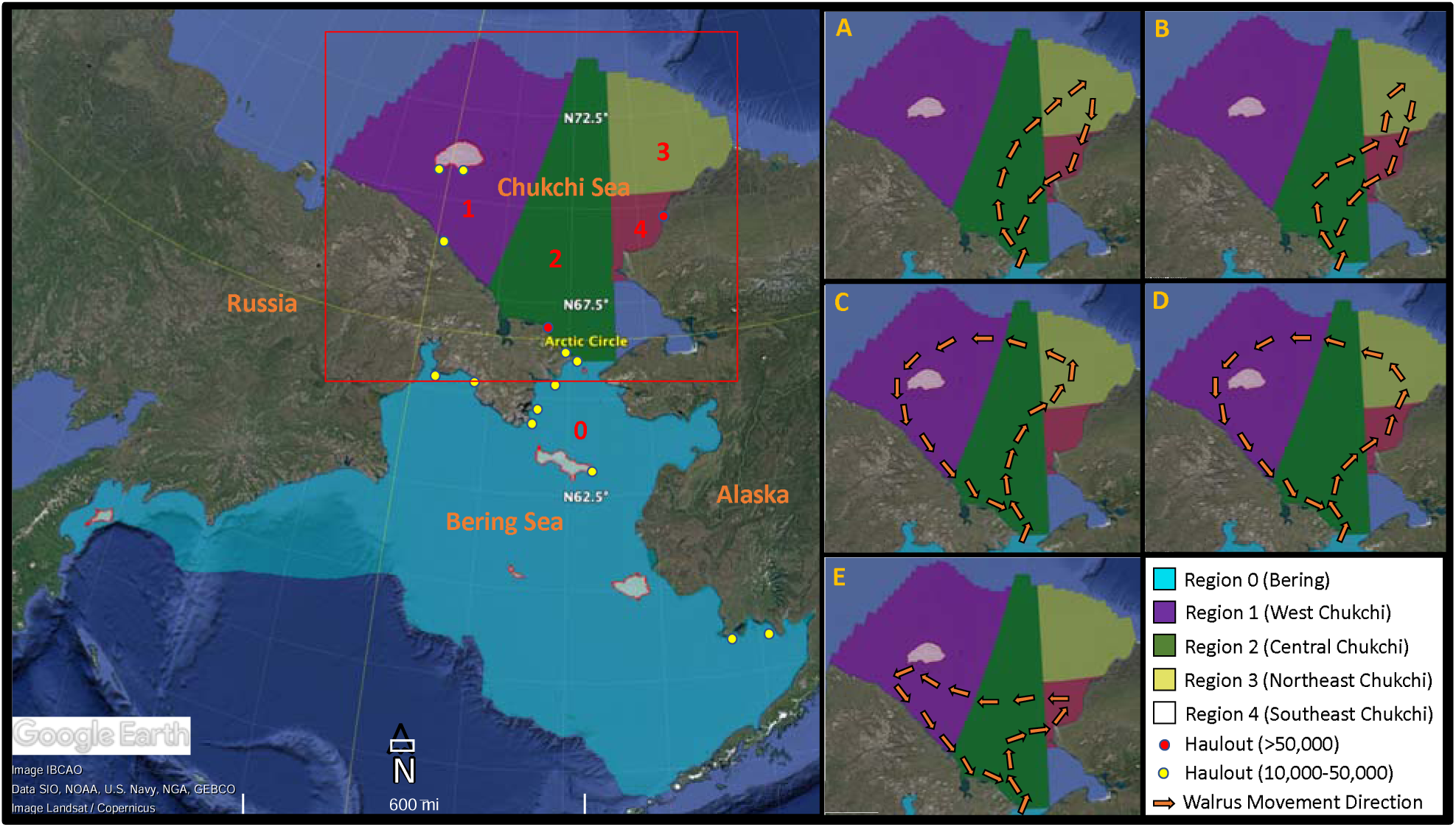
Map of the study area, denoting the annual range of the Pacific walrus divided into the five regions (0-4) considered in this study. The Kotzebue Sound portion of the Chukchi Sea (south of Region 4 and east of Region 2) is not walrus habitat and was not included in the study. Smaller panels (A-E) represent the five movement pathways during the summer period in the Chukchi Sea (June-November) identified by Udevitz et al., (2017) and used in the dynamic energy budget. Gray shaded regions are islands. Points along the coasts correspond to routinely-used coastal haulout sites with aggregations of >10,000 – 50,000 and >50,000 individuals (MacCracken et al., 2017; Fischbach et al., 2022).

The CMIP6-endorsed global coupled ocean-atmosphere general circulation models (Fox-Kemper et al., 2021) we used to project sea ice cover did not contain a set of models directly analogous to the CMIP5 models used by Udevitz et al. 2017; thus, we initially considered a set of 13 models identified by the SIMIP (Sea Ice Model Intercomparison Project) Community (2020) to best estimate the future evolution of Arctic/subarctic sea ice cover. We excluded three models that did not have data available for both the intermediate (ssp245) and pessimistic (ssp585) scenarios, resulting in a set of ten models (Table 2). For each climate model, we considered two 10-year time periods (mid-century = 2045–2054; end-century = 2090–2099), and two shared socio-economic pathways (ssp245 & ssp585). We applied a sampling protocol to incorporate the variability between models, resulting in a total of five sea ice scenarios (SI_0– SI_4; Table 2; Fig. 3).

**Figure 3.**
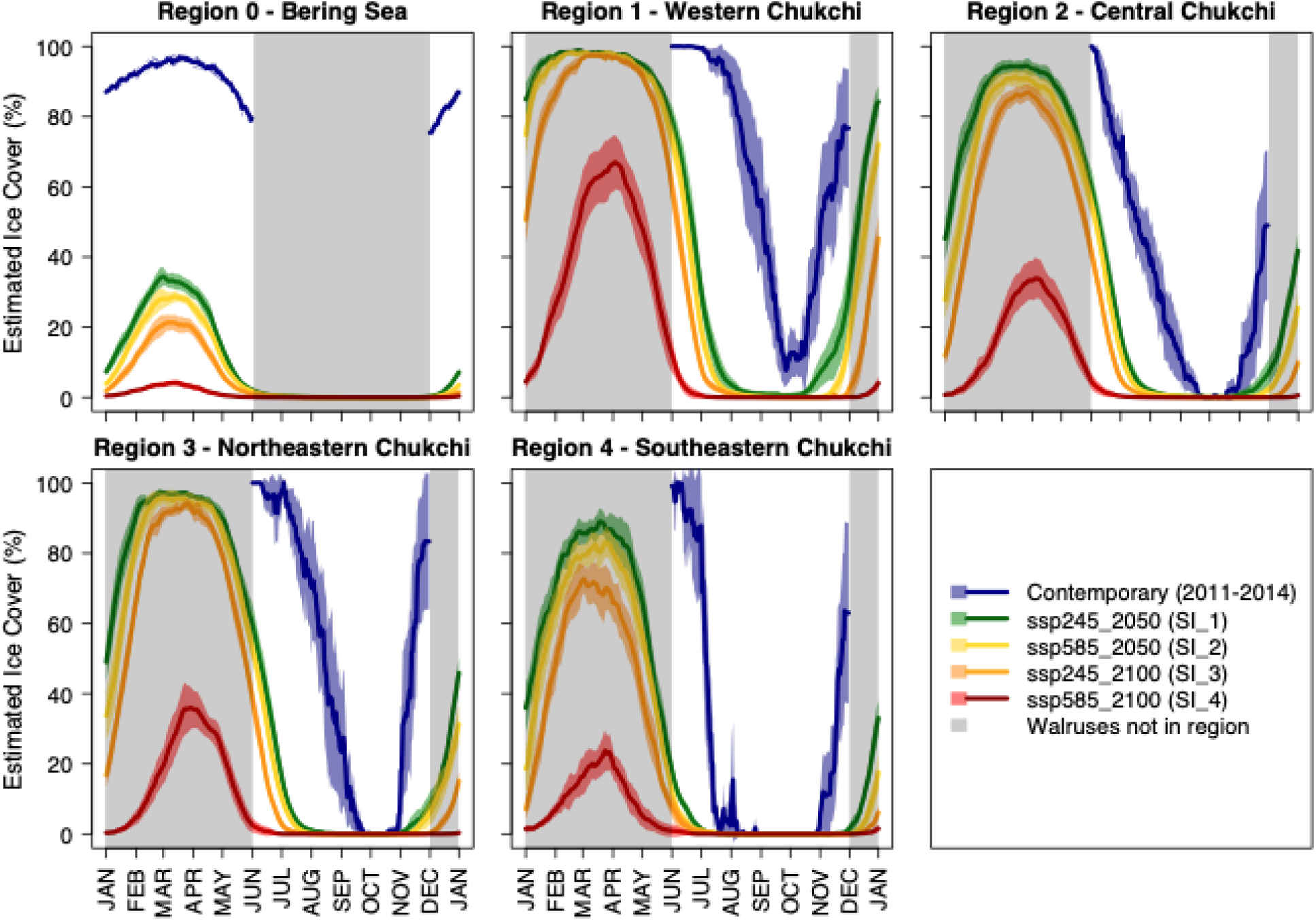
Sea ice projections for the five regions of the study area, based on historical data (blue) and four climate scenarios (green, yellow, orange, red) averaged across the suite of ten CMIP-6 climate models used in this study. For ssp245_2050 & ssp585_2050, sea ice projections include projections from 2045-2054, and for ssp245_2100 & ssp585_2100, sea ice projections include projections from 2090-2099. Solid lines represent the mean of 10 years of projected data for each day of the year, and shaded polygons are the 95% density intervals representing the variability across years and models. Gray rectangles represent periods of time that modelled walruses are not in their respective regions (i.e., the Chukchi Sea during the winter and the Bering Sea during the summer). Contemporary estimates of sea ice were not available for non-walrus periods.

**Table 2.** The five sea ice scenarios used to assess population consequences of climate change. Shared Socio-Economic Pathways (SSPs) were developed by the IPCC (International Panel on Climate Change) and represent an intermediate (ssp245) and pessimistic (ssp585) carbon emission mitigation scenario.

| Name | SSP | Projection Period | Ice Data Source |
| --- | --- | --- | --- |
| SI_0 | NA | 2008–2014 (contemporary) | MIZ & SIGRID-3<sup>1</sup> |
| SI_1 | ssp245 | 2045–2054 (mid-century) | CMIP6 Suite<sup>2</sup> |
| SI_2 | ssp245 | 2090–2099 (end-century) | CMIP6 Suite<sup>2</sup> |
| SI_3 | ssp585 | 2045–2054 (mid-century) | CMIP6 Suite<sup>2</sup> |
| SI_4 | ssp585 | 2090–2099 (end-century) | CMIP6 Suite<sup>2</sup> |

<sup>1</sup>Historical sea ice data for the 2008–2014 period were compiled by Udevitz et al. (2017) using MIZ (Marginal Ice Zone) and SIGRID-3 (Sea Ice Grid 3) charts.
<sup>2</sup>For each sea ice projection, we applied a sampling protocol to randomly incorporate data from the following suite of CMIP6-endorsed models (institute and host country): ACCESS-CM2 (CSIRO-ARCCSS, Australia), ACCESS-ESM1.5 (CSIRO, Australia), CanESM5 (CCCma, Canada), CESM2-WACCM (NCAR, U.S.A), CNRM-ESM2-1 (CNRM-CERFACS, France), EC-Earth3 (EC-Earth Consortium, Europe-wide), EC-Earth3-Veg (EC-Earth Consortium, Europe-wide), IPSL-CM6A-LR (IPSL, France), MIROC6 (MIROC, Japan), MRI-ESM2-0 (MRI, Japan).

We defined the female Pacific walrus wintering range (from December–May) as the maximum sea ice extent (including the marginal ice zone as well as pack ice) of the Bering Sea (Fig. 2) which corresponds to previous estimates of the Pacific walrus range (e.g., MacCracken et al., 2017). Because little is known about Pacific walrus winter movements, we considered this portion of the Bering Sea as one region within our analysis. To characterize contemporary ice cover in the Bering Sea region, we applied the protocol outlined in Udevitz et al., (2017), analyzing MIZ & SIGIRD-3 ice charts for December–May 2008–2015. From the Chukchi Sea dataset, we generated distributions of the response of Pacific walruses (across five movement pathways; Udevitz et al., 2017) to varying degrees of ice cover and applied them to Bering Sea ice cover estimates to simulate walrus behavior in the Bering Sea.

Finally, we developed six movement scenarios based on the five summer movement patterns from Udevitz et al. (2017). Our first five scenarios (one for each movement pattern) assumed that each female’s movement pattern remained consistent throughout her lifetime, but we also include a sixth scenario where the female’s movement pattern was resampled annually. Because the proportion of the population that follows each movement pattern in any year is unknown, and year to year fidelity to patterns is also unknown, we assigned an equal probability of occurrence to each of the six movement scenarios and conducted a sensitivity analysis to these scenarios.

We used both the sea-ice-driven movement and activity models from Udevitz et al. (2017), to generate Bayesian posterior predictive distributions of daily activity budgets for Pacific walruses for each movement scenario under the five different sea ice scenarios (Table 2). The samples from these distributions generated daily estimates of the following activity budget parameters: the percentage of time an individual spends in the water (*P_WATER_*); the percentage of time it spends foraging (*P_FORAGE_*); and the region it occupies on each day. These values link the daily field metabolic cost (*FM_t_;* Equation 9) and intake rate (*IR_t_*; Equation 3) to ice cover. Our baseline model uses sea ice estimates for contemporary conditions (i.e., SI_0; Table 2), and models incorporating sea ice projections (SI_1–4) simulate the response of Pacific walrus energy intake and expenditure to ice loss.

Female walruses and their calves have traditionally used ice during the summer/autumn period in the Chukchi Sea as a platform to rest on between foraging bouts, which provides easy access to foraging areas and a refuge from terrestrial predators and anthropogenic disturbance (Fay 1982). As the summer/autumn ice-free period has increased, larger numbers of walruses have used terrestrial haulouts instead (Fischbach et al., 2009; Fischbach et al., 2022), which increases the travel time to foraging grounds (Jay et al., 2017) and the probability of disturbance-related mortalities (Udevitz et al., 2013). Young calves are particularly susceptible to mortality at terrestrial haulouts, both from predators (e.g., polar bears), and from being trampled in disturbance-initiated stampedes (Fischbach et al., 2009). Terrestrial haulouts on the Chukchi coast have been monitored in recent years and can be used by >150,000 individuals in some instances (Fischbach et al., 2022), and minimum annual haulout mortality estimates ranged from 20–3400 individuals between 2007 and 2016 (MacCracken et al., 2017). Pacific walrus haulout mortality may increase as a function of sea ice loss (Udevitz et al., 2013).

To account for haulout mortality in our climate scenarios, we created a terrestrial haulout day parameter (*THD*) which we defined as:

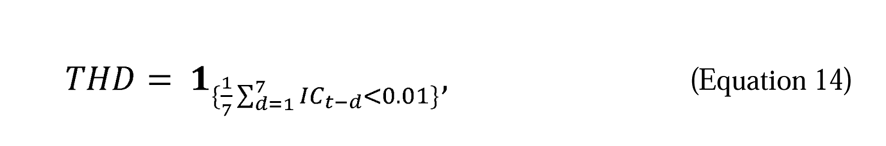

where *IC* is the ice cover in the simulated individual’s current region. In other words, if an individual has been in a region with a mean of <1% ice cover over the past 7 days, it must rest at and base from a terrestrial haulout. We base this estimate of 7 days on observations of walrus behavior at sea (e.g., in Fay 1982), and on preliminary analysis of satellite tracking data paired with ice availability (Fischbach & Jay, 2018). Under the baseline model, an individual has an average of 40 *THD*s (Fig. S9), which is consistent with contemporary estimates (Fischbach & Douglas, 2022). Note that each *THD* is simply a day a simulated walrus spends basing from a terrestrial haulout (rather than ice), and it does not confer 24 hours spent hauled out on land. We used haulout mortality estimates from 2007–2016 (MacCracken et al., 2017) alongside genetic mark-recapture population size estimates (Beatty et al., 2022) and demographic haulout mortality data from both Alaska and Russia (Fischbach et al., 2016) to estimate the annual probability that an adult female or her calf dies at a terrestrial haulout. These estimates were divided by the number of *THD*s under contemporary conditions (37) to attain the daily probability of terrestrial haulout mortality (,*_THM_F_* for adult females and,*_THM_C_* for calves; Table S3). Each,*_THM_* parameter was calculated and applied separately for Alaska and Russia, and the mean of the Alaska and Russian values was used for the Bering Sea (Region 0) because it included terrestrial haulouts in both countries. On each *THD*, we performed a Bernoulli trial with success probability,*_THM_C_* to determine calf survival and,*_THM_F_* to determine adult survival.

In addition to the baseline,*_THM_* parameters, we included “bad year” (*BY*) parameters to simulate the natural stochasticity that has been observed with large-scale haulout mortality events (e.g., Udevitz et al., 2013). Large-scale haulout mortality events occur periodically when females and their calves are disturbed when occupying terrestrial haulouts in particularly high densities, resulting in mass mortalities from stampeding (e.g., Ovsyanikov et al., 2007; Fischbach et al., 2009). To simulate *BY*s in the model, we estimated the daily probability of a calf or adult dying at a terrestrial haulout in 2007 (the only historical “bad year” for which we have reliable data) and applied that value to the Bernoulli trial on each *THD* (probabilities,*_THM_BY_F_* for adult females and,*_THM_BY_C_* for calves) on X randomly-selected “bad years” of a female walrus’s lifespan.

Finally, we added a haulout mortality management (*HMM*) parameter to simulate the effect of management and conservation efforts on mitigating coastal haulout mortalities. This parameter (ranging from 0.0–1.0) is multiplied by one minus the probability of calf haulout mortality (,*_THM_C_*) or adult haulout mortality (,*_THM_F_*), ultimately reducing it. For instance, an *HMM* value of 0.2 would reduce the baseline haulout mortality probability by 20%, simulating management/conservation efforts such as maintaining a buffer zone around known terrestrial haulouts to reduce disturbance (e.g., Garlich-Miller et al., 2011). The probability of calf or adult haulout mortality in “bad years” (,*_THM_BY_C_* or,*_THM_BY_F_*), was not modified by the *HMM* parameter to maintain the simulated stochasticity of such events.

Thus, an individual’s survival in any given scenario was determined by the product of 2–4 Bernoulli trials, in the following order, to determine if the individual survived (1) the background mortality rate based on its age at death under good conditions, (2) starvation (if applicable), (3) terrestrial haulout-based mortality (which may or may not have been mitigated by management), and (4) the effects of a bad terrestrial haulout year (if applicable).

Little is known about the composition and biomass of benthic invertebrates in the Chukchi Sea and how changing sea ice availability, warming subsurface sea temperatures, phytoplankton blooms (Arrigo & van Dijken, 2015), and range expansions of pelagic species (e.g., Huntington et al., 2020) may ultimately influence Pacific walrus prey density (MacCracken et al., 2017). There is some evidence from focused regional studies that indicates a decline in benthic biomass in the southern Chukchi Sea (Grebmeier et al., 2015), but, in contrast, a longer ice-free interval could theoretically increase foraging opportunities, as has been postulated for Atlantic walruses (*Odobenus rosmarus rosmarus*; Laidre et al., 2008). To represent changes to prey density in projected climate change scenarios, we used a parameter (*PD*) which proportionally weights the resource availability parameter (*R*; Table 1).

### POPULATION CONSEQUENCES OF DISTURBANCE

Several different types of anthropogenic disturbance may adversely impact walrus behavior (Table S1; MacCracken et al., 2017), but little information exists on specific behavioral responses that walruses may exhibit. An expert elicitation (EE) was conducted in 2019 to generate expert opinion-based probability distributions of transfer functions from acoustic anthropogenic disturbance to Pacific walrus behavior (Harwood et al., 2019). The EE quantified the reduction of an adult female Pacific walrus’ daily foraging effort in response to an acute acoustic stressor (i.e., a seismic survey) and to a continuous acoustic stressor (i.e., a drilling operation; Fig. S10).

The DEB framework uses these transfer functions to link behavioral responses to disturbance, and ultimately to population-level effects. Daily behavioral responses that result from disturbance can be incorporated into scenarios by specifying the following parameters: disturbance type (seismic or drilling), the number of disturbance days/year, whether those disturbance days occur randomly during the June-November period or during a specified timeframe, whether disturbance occurs during consecutive days or years, the number of years in an individual’s lifetime that disturbance takes place, and whether those disturbance years occur randomly or at a specified age. This framework allows managers the flexibility to simulate specific development projects (and their effects on different age classes), or to consider disturbance under more generalized scenarios.

### SCENARIOS

Based on input from wildlife managers and IK stakeholder groups, we identified four primary climate/disturbance scenarios to demonstrate our modeling framework (Fig. 4). Although we include and discuss these four scenarios, the PCoD framework is highly flexible and designed to assess wide array of additional scenarios. Each scenario incorporates the following variables: a climate scenario (intermediate: ssp245 or pessimistic: ssp585) and the associated effects of sea ice availability; changes to prey density (*PD*); the haulout mortality management (*HMM*) factor; the number of “bad years” for haulout mortality (BY); and the amount of anthropogenic disturbance (e.g., from drilling or seismic surveys). We conducted sensitivity tests to assess the individual effects of these variables on model output, and ultimately to determine their relative importance in the four combined scenarios. In our scenarios, we considered disturbance days to occur randomly in randomized years. Each disturbance day had a 50% chance of the disturbance being seismic vs. drilling, and the simulated individual’s foraging intake was modified by a random draw from the distribution associated with each disturbance type (Fig. S10). For the range of variation considered for each variable (e.g., 1–3%, Fig. 4) we selected a random value for each simulated individual. For each of the four scenarios (Optimistic, Intermediate_ssp245 Intermediate_ssp585, Pessimistic; Fig. 4), we estimated *K* and *r_max_* for the years 2050 and 2100, each with 100 simulations each comprising 100 individuals.

**Figure 4.**
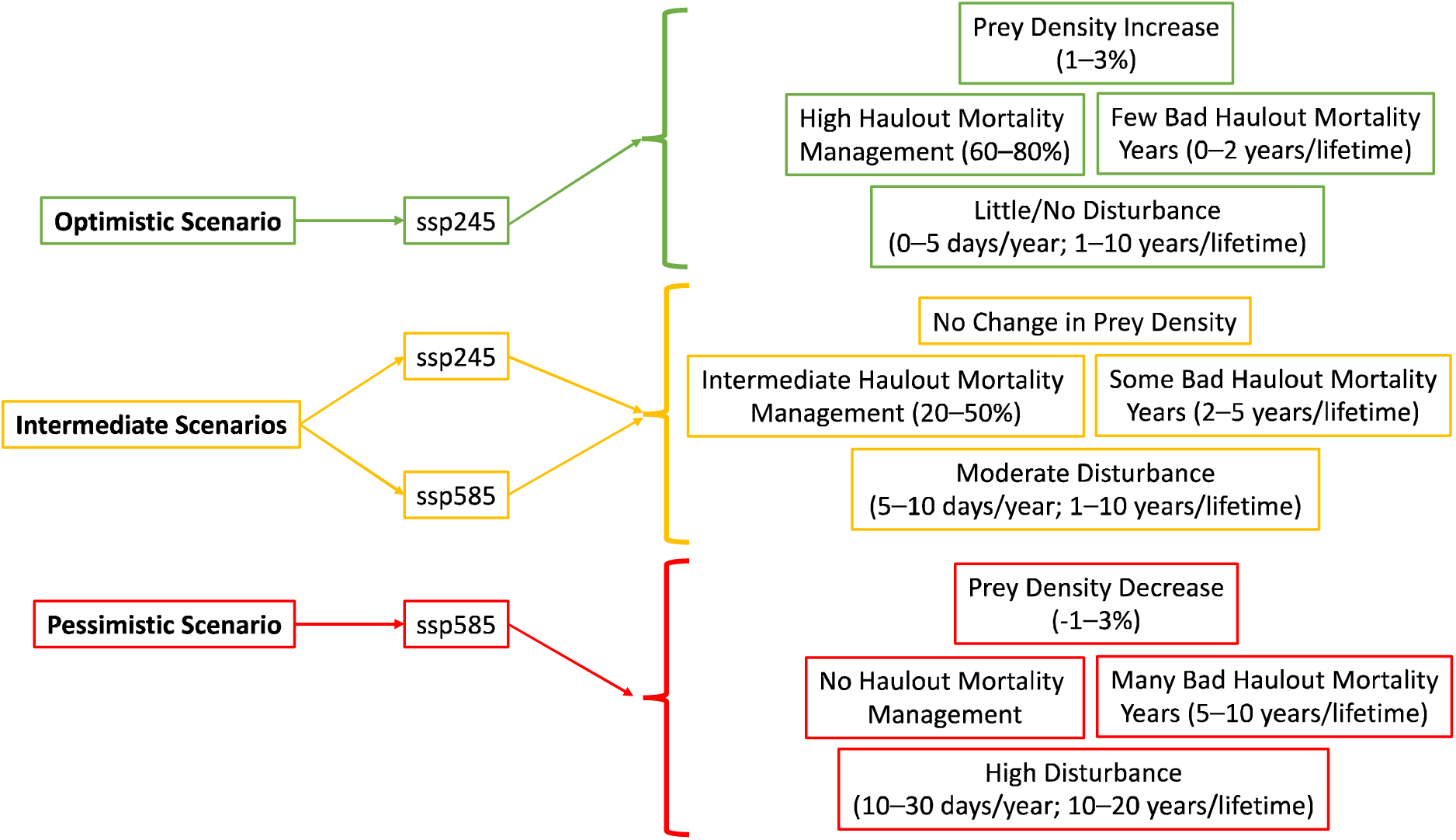
Summary of climate/disturbance scenarios developed for the present study. ssp245 (intermediate) and ssp585 (pessimistic) refer to IPCC (Intergovernmental Panel on Climate Change) CO_2_ emissions scenarios and their associated sea ice projections.

### MODEL SENSITIVITY

We conducted a series of analyses to assess the model’s sensitivity to several core bioenergetic parameters and scenario parameters. These included the target body condition threshold (ρ), the starvation threshold (ρ*_*s), prey density (*PD*), and anthropogenic disturbance (both days/year and years/lifetime). Statistical significance between groups of model parameters was determined via ANOVA and post-hoc Tukey tests. All analyses were conducted using the R statistical software (R Core Team, 2023).

## RESULTS

### Calibration of Dynamic Energy Budget

The Dynamic Energy Budget was effectively calibrated to reproduce demographic rate estimates for 2015 produced via an IPM (Taylor et al., 2018 most parsimonious model; Table S2). Specifically, ≥70% of simulations calibrated to an *R* and φ*_L_* of 4.034 (n = 200 simulations each containing 100 individuals) fell within a 95% CrI of IPM estimates: 70% for reproduction, 100% for neonatal survival, and 100% for older calf survival (Fig. S7). Under these conditions, the simulated population’s *N/K* value approximates 0.9 when ρ = 0.3 and ρ_s = 0.1 (i.e., the values in Table 1), although the model was not shown to be sensitive to those body condition thresholds (Fig. S11). Thus, we used the DEB calibrated to this *R* and φ*_L_* under contemporary conditions (the SI_0 sea ice scenario; Table 2) to serve as the baseline model in our PCoD framework. Figure 5 shows an example of one female walrus’s simulated mass balance and reproductive success under this model.

**Figure 5.**
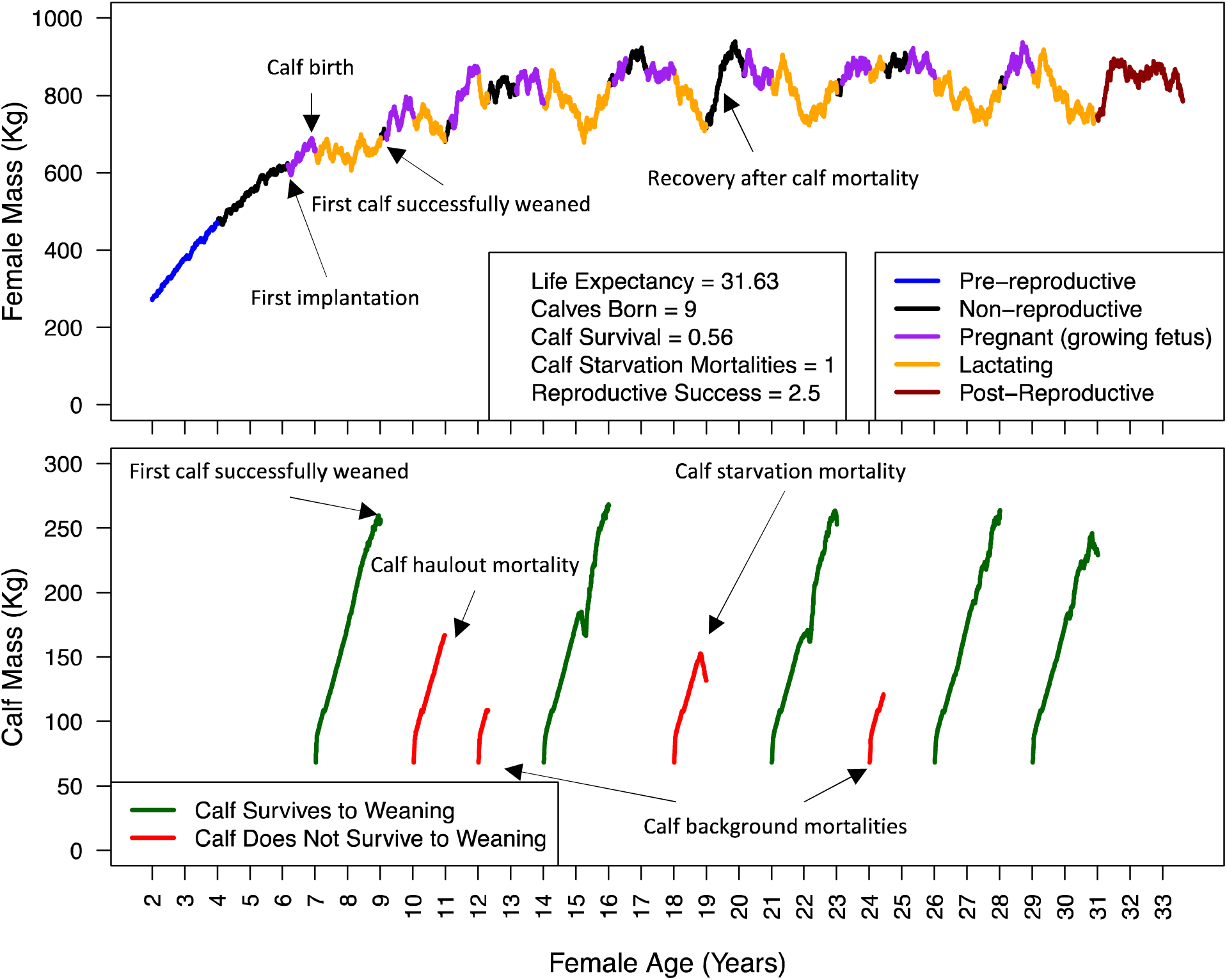
Example output for one simulated female walrus over the course of the simulation. The top panel reflects fluctuations in female body mass associated with different activity states, and the bottom panel represents concurrent reproductive attempts over the female’s lifetime. Reproduction refers to the total number of female calves this walrus successfully reared during her lifetime (total calves/2).

### Sea ice Projections

CMIP6 sea ice projections indicated significant declines in ice cover across the five study regions in all four scenarios (Fig. 3). Generally, this corresponds to a months-long ice-free interval across all Chukchi Sea regions during the summer/autumn months, and reduced sea ice availability during the traditional walrus breeding season in the Bering Sea. Most notably, the pessimistic (SI_4) sea ice scenario suggests that the traditional annual range of the Pacific walrus will be almost entirely ice-free by the end of the 21^st^ century (Fig. 3).

### Walrus Response to Climate-Induced Changes

Simulated walruses had higher energy expenditure with reduced ice cover in all climate scenarios, which resulted from their need to spend a higher proportion of time active in the water (Udevitz et al., 2017). With all other factors held constant in the model, this sea ice-induced increase in energy expenditure significantly impacted starvation-related mortality rates and reproductive success. Specifically, populations modelled under the SI_1, SI_2, and SI_3 scenarios had significantly higher rates of both adult and calf starvation, and lower calf survival (probability of the calf surviving to weaning at 2 years old) and reproductive success than the SI_0 (baseline) scenario (Fig. S12). The SI_4 scenario (representing the ssp585 climate scenario at the end of the 21^st^ century) exhibited the highest rates of adult and calf starvation, and lowest rates of reproductive success and calf survival (Fig. S12).

Similarly, the probability a simulated walrus used a terrestrial haulout increased above the 2015 mean of 37 THDs under all climate scenarios. Under the pessimistic (ssp585) scenario, walruses experienced a mean of 96 THDs in 2050, and 248 THDs in 2100 (Fig. S9). When projecting forward to 2100, we predicted a higher proportion of THDs in the Russian Chukchi Sea and the Bering Sea than in the Alaskan Chukchi Sea, suggesting that the regional distribution of terrestrial haulouts will also change. Increased time at terrestrial haulouts created an additional source of calf and adult mortality in our climate scenarios which contributed to the decline in reproductive success (Fig. S13). The model was relatively sensitive to “bad years” for terrestrial haulout mortality (BYs). Adding BYs significantly increased both calf and adult haulout mortality in an incremental fashion, which contributed to a significant decline in reproductive success when each simulated individual was subjected to 20 bad haulout years in her lifetime (Fig. S13).

The DEB was highly sensitive to the prey density (*PD*) scaling parameter which scaled *R* (resource availability). Although a baseline DEB calibrated to an *R* of 4.034 produced demographic rates consistent with empirical estimates for the population (e.g., Fig. S7), adjusting that *R* by ± 1–5% had large impacts on starvation rates and reproductive success (Fig. S14).

Finally, we conducted an analysis to assess the effect of sea ice scenarios on the five walrus movement patterns (A–E). We found significant differences in adult/calf starvation, calf survival, and associated reproductive success between the movement patterns (Fig. S15). Specifically, Movement Pattern A (one of two patterns confined to the eastern Chukchi Sea) conferred the lowest instances of starvation, and highest calf survival and reproductive success, whereas Movement Pattern E (one of three patterns using the eastern and western Chukchi Sea, but with the least northerly extent) conferred the highest instances of starvation and lowest calf survival and reproductive success, and the other three movement patterns fell between the two. These differences remained relatively consistent regardless of which sea ice scenario was applied (Fig. S15).

### Walrus Response to Anthropogenic Disturbance

Simulated walrus demographic rates were moderately sensitive to anthropogenic disturbance. When seismic or drilling disturbance occurred at random days during the year on 5 or 10 random years over a simulated walrus’ lifetime, an extremely large number of disturbance days (i.e., 30) per year was required to significantly impact calf or adult starvation rates and associated reproductive success (Fig. S16). Exposure to 10 disturbance days for 20 years/lifetime was enough to significantly reduce reproductive success, and the most extreme disturbance scenario (30 days/year for 20 years/lifetime) reduced overall reproductive success by 45% (Fig. S16). Although we considered anthropogenic disturbance in a randomized fashion in the present analyses, disturbance may realistically have a greater impact on starvation rates and reproductive success if it: A) occurs on consecutive days/years; B) occurs in a localized critical habitat area (e.g., an important seasonal foraging ground); or C) occurs during a time of year when walruses are more vulnerable (e.g., just after calving).

### Combined Climate/Disturbance Scenarios

The four primary scenarios combining climate change and disturbance (Fig. 4) all indicated declines to both Pacific walrus carrying capacity (*K*) and intrinsic rate of increase (*r_max_*) by the end of the 21^st^ century, and sooner in most cases (Fig. 6). Under the most optimistic scenario, *K* underwent a gradual decline (95% CrI for 2015: 78106–82060; for 2100: 64037–67875), as did *r_max_* (95% CrI for 2015: 0.038–0.039; for 2100: 0.032–0.033). The Intermediate_245 model resulted in a slightly stronger decline in *r_max_* to the optimistic scenario (95% CrI for 2100: 0.028–0.029), and a larger, quicker decline in *K* (95% CrI for 2100: 46144–48249). The Intermediate_585 model resulted in a stronger decline in K than the Intermediate_245 model (95% CrI for 2100: 37988–39546), matched with a stronger decline in *r_max_* (95% CrI for 2100: 0.023–0.025). Finally, our pessimistic scenario indicated severe declines to both *K* (95% CrI for 2100: 25715–26724) and *r_max_* (95% CrI for 2100: 0.017–0.019; Fig. 6). Ultimately, this pessimistic scenario would amount to a 67% decline in carrying capacity and a 53% decline in *r_max_* by the end of the 21^st^ century.

**Figure 6.**
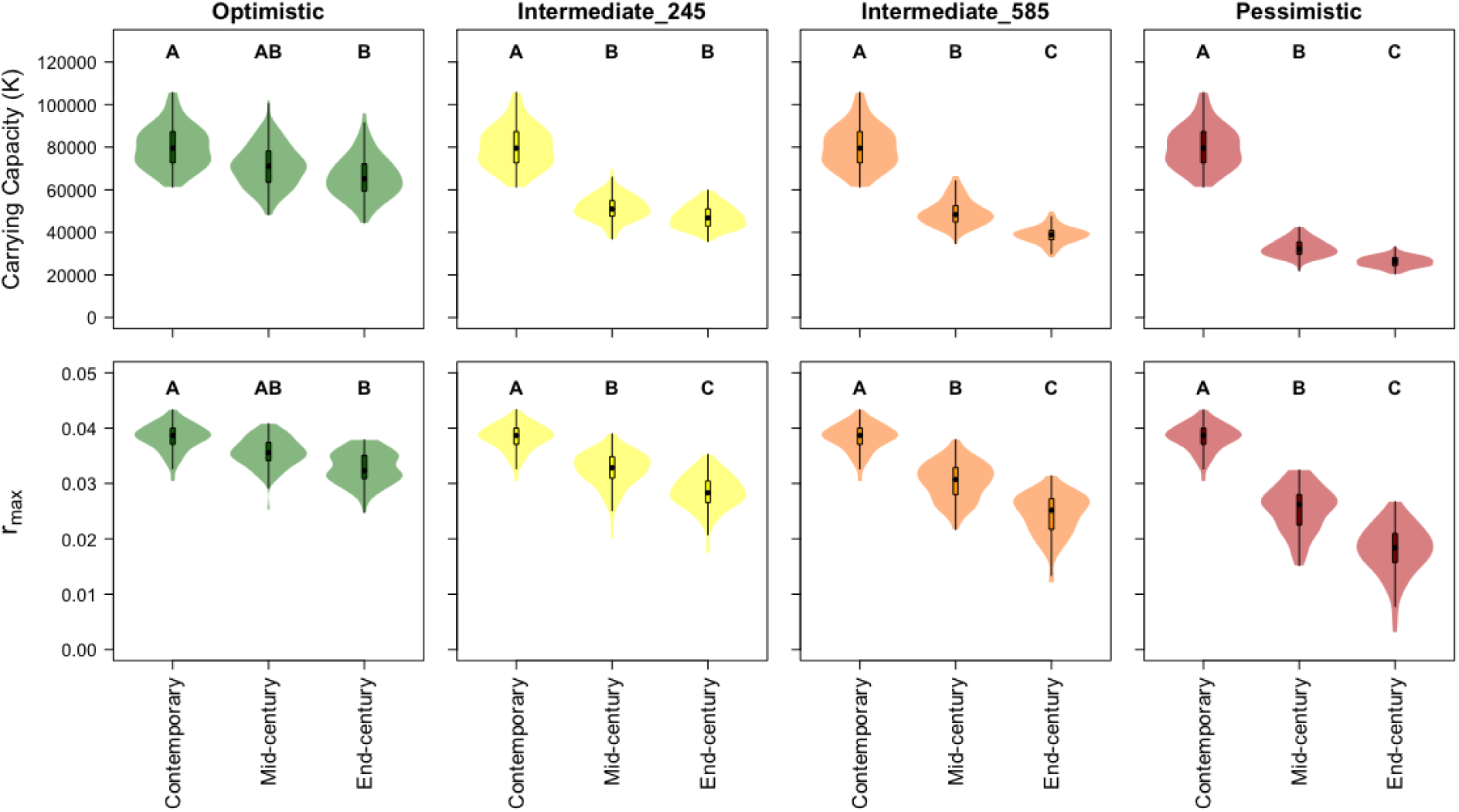
Projected *K* (carrying capacity; top panels) and *r_max_* (intrinsic maximum rate of population growth; bottom panels) estimates for the adult female Pacific walrus population to the middle and end of the 21^st^ century under four combined disturbance/climate change scenarios in the PCoD framework (described in Figure 4). Contemporary estimates incorporate sea ice conditions between 2008-2015; mid-century estimates 2045-2054; end-century estimates 2090-2099. Different letters indicate significant differences between groups from an ANOVA and post-hoc Tukey test.

## DISCUSSION

The PCoD model provided a useful framework framework to analyze consequences to the Pacific walrus population of climate change-induced sea ice loss—a well-recognized and growing concern among the scientific community (e.g., MacCracken 2012; Fischbach et al., 2022), indigenous communities that rely on walruses for subsistence, and the general public. The model allows for direct and indirect effects of sea ice loss on individual behavior to be translated into changes in population demography and size. A central strength of our PCoD model is that it can incorporate new information as it becomes available, and scenarios can be tailored to evaluate the stressors of greatest concern.

We evaluated four example scenarios (Fig. 4), all of which resulted in a statistical decline in both carrying capacity (K) and the intrinsic rate of increase (r_max_), and thus overall abundance and population growth rate to the end of the 21st century. Projected changes to the population were primarily driven by the rate of sea ice loss, mortality at terrestrial haulouts, and changes to prey density. Diminishing sea ice resulted in walruses swimming more and foraging less, thus increasing energy expenditure, and decreasing energy intake. This increased adult and calf starvation rates, and reduced reproductive success and calf survival (Fig. S12), which, in turn, drove the decrease in carrying capacity (Fig. 6). Reduced prey density and increased anthropogenic disturbance similarly decreased the daily walrus energy intake rate, reducing carrying capacity. The DEB was highly sensitive to prey density and relatively sensitive to anthropogenic disturbance. Our variables associated with terrestrial haulout mortality (the number of bad haulout years and the strength of the haulout management effect) were modeled independently of resource availability and were thus the primary drivers of the observed trends in rmax.

Movement patterns are a critical aspect of our model, and these patterns were based on data from telemetered walruses and the timing of their movements among regions relative to the amount of regional sea ice (Udevitz et al., 2017). We note, however, that the overwhelming majority of animals tracked by Udevitz et al. (2017) were satellite-tagged in the U.S. portion of the Chukchi Sea, and in the absence of any additional information, we assumed these movement patterns (Fig. 2, patterns A–E) represented all major movement patterns exhibited by the population and that each pattern was equally likely (i.e., each has a 1/5 probability of being assigned to an individual). We also assumed the direction of these movement patterns will remain consistent up to the end of the 21st century. Despite these limitations, the strength of our movement pattern modeling is that residence within any given region in any movement pattern is linked to the amount of sea ice in that region; thus, our model accounts for a shift in migratory phenology as a result of sea ice loss (Udevitz et al., 2017).

Although our PCoD framework has the capacity to incorporate many of the previously identified potential stressors to the Pacific walrus population (i.e., Table S1), further study is required to encompass the full suite of possible stressors associated with climate change and anthropogenic disturbance. For example, impacts of Bering Sea ice loss on breeding and birthing platforms may have ramifications on reproductive success that are not accounted for in the model due to lack of empirical data. Pacific walruses are thought to require sea ice platforms to breed from and give birth on (MacCracken et al., 2017), so if an ice-free Bering Sea (e.g., in the SI_4 scenario) did not result in a range shift (e.g., a shift northward to sea ice refugia resulting in year-round residence in the Chukchi Sea, MacCracken 2012), our model does not account for the potential shift from breeding from and birthing on ice to breeding from and birthing on land, which could result in large reductions in reproductive success. Similarly, adult female walruses are not known to leave their calves on land and forage alone at sea (as do sea lions, *Otariidae*), thus we assumed that walrus calves are able to accompany their mothers everywhere, and that they have the in-water endurance to do so in an ice-free environment. In addition, further information is needed on effects of climate change to the benthic community (and how that relates to future walrus prey density; but see Grebmeier et al., 2015; Wilt et al., 2013) and prevalence of harmful algal blooms and other diseases and parasites. An expert elicitation (EE) comprised of agency walrus biologists produced the probabilities our PCoD model used to quantify walrus responses to seismic and drilling disturbance (Fig. S10, Harwood et al., 2019). An additional EE (that includes IK stakeholders) could prove beneficial for acquiring similar expert opinions for other disturbance types (e.g., ship and air traffic, fisheries, and military exercises) as well as for filling additional knowledge gaps with regards to Pacific walrus physiological and behavioral ecology.

Most bioenergetic models are, by necessity, simplifications of extremely complex natural processes. Our DEB is based on relationships among parameters that incorporate empirical values from Pacific and Atlantic walruses, assumed values and relationships based on walrus life history, and data or assumptions from other marine mammals where walrus-specific data were not available (Table 1). For instance, we applied a starvation threshold from a study on Steller sea lions (*Eumetopias jubatus*; Noren et al., 2009), and several of our scaling and shape parameters were drawn from the DEB presented by Hin et al. (2019) who chose them to best reflect assumed biological relationships for long-finned pilot whales (*Globicephala melas*). As such, we have identified several data gaps in the Pacific walrus and marine mammal bioenergetic literature that had been filled with assumed values in the present model. Key research needs related to Pacific walrus physiology include activity-associated metabolic rates for growing calves, further research on walrus diet and prey density in the Bering and Chukchi seas, and a more robust characterization of assimilation efficiency, true metabolizable energy, and the heat increment of feeding (e.g., Booth et al., 2023). Despite the necessary assumptions we have made within this modeling framework regarding bioenergetics, its strength lies in its flexibility to compare and contrast different parameter values and projected environmental conditions.

The PCoD model is a flexible framework that should prove useful to wildlife managers and stakeholders to project and assess walrus population dynamics under a range of potential future conditions. In addition, while current subsistence harvest rates are thought to be sustainable (USFWS 2023), a formal harvest sustainability assessment has not yet been conducted for the population (MacCracken et al., 2017), and it is an active area of research. Estimates of K and r_max_ attained from our PCoD model could be used to form a baseline for a harvest sustainability assessment which can project the sustainability of different harvest scenarios to the end of the 21^st^ century. Finally, the framework we have developed could be readily adapted to other situations and species in complex, dynamic systems. It will be particularly useful for the many species simultaneously threatened by climate change and anthropogenic disturbance.

## ACKNOWLEDGEMENTS

We are extremely thankful to members of the Pacific Walrus Harvest Model Steering Committee and the Eskimo Walrus Commission for their support and discussion in the development of this modeling framework: Charles Brower, Vera Metcalf, Jacob Martin, Bryan Rookuk Jr., and Enoch Oktollik. We are also grateful to John Harwood and Cormac Booth for developing an interim Pacific walrus DEB model upon which this analysis is based. Erik Anderson, Jen Cate, and Patrick Lemons provided useful input on model development and thoughtful review to a previous version of this manuscript. The findings and conclusions in this article are those of the author(s) and do not necessarily represent the views of the U.S. Fish and Wildlife Service. This product paper has been peer reviewed and approved for publication consistent with USGS Fundamental Science Practices (https://pubs.usgs.gov/circ/1367/). Any use of trade, firm or product names is for descriptive purposes only and does not imply endorsement by the U.S. Government.

## CONFLICT OF INTEREST STATEMENT

The authors declare no conflict of interest.

## Notes

### Competing Interest Statement

The authors have declared no competing interest.

