## Supplemental Tables & Figures for "Assessing the Population Consequences of Disturbance and Climate Change for the Pacific Walrus"

Table S1. Primary potential stressors to the Pacific walrus population [updated from MacCracken et al., 2017]. Stressors not currently (or partially) encompassed in model scenarios require additional data or expert opinion in order to be incorporated.

| **Potential Stressor** | **Potential Effect on Pacific Walruses** | **Relative Intensity of Stressor in 2100** | **Encompassed by model** | **Data-based inclusion in scenarios** | **Expert-opinion based inclusion in scenarios** | **Inclusion in scenarios but more information needed** | **Data gaps** |
| --- | --- | --- | --- | --- | --- | --- | --- |
| CLIMATE STRESSORS |  |  |  |  |  |  |  |
| Loss of summer/fall ice in Chukchi Sea | Increased energy expenditure (more time active in water, not foraging); Increased coastal crowding | High | Yes (*P_WATER_*, *P_FORAGE_*, *THD*) | Sea ice projections, effect on foraging, terrestrial haulout mortality | N/A | N/A | Updated data regarding walrus movement and activity budgets with regards to coastal haulout use could be incorporated when it becomes available. |
| Decline in winter/spring sea ice in Chukchi and Bering Seas | Northward shift in distribution;  Decline in sea ice breeding platforms (winter) & birthing platforms (spring) | High^1^ | Yes (*P_WATER_*, *P_FORAGE_*, *THD*) | N/A | Sea ice projections, effect on foraging, terrestrial haulout mortality | N/A | Data on activity budgets and terrestrial haulout mortality were collected primarily in the Chukchi Sea in summer/autumn. No information about concerns in column 2. |
| Climate change effect on benthic prey (ocean warming & acidification) | Changes in prey abundance, distributions, and species composition | Moderately High - High | Yes (*R*, *PD*) | N/A | N/A | Yes, through prey density, but not mechanistically | Data needed before mechanistic modeling can be done; Comprehensive surveys of walrus forage species across the Chukchi/Bering seas. |
| Disease & Parasites | Increased likelihood of disease and parasites due to increased haulout crowding | Low-Moderately Low | Yes ($\phi$*_THM_*_)_ | N/A | N/A | Yes, through increased haulout mortality, but not modeled mechanistically | Data needed before mechanistic modeling can be done; little data exist regarding walrus disease or parasites. |
| Predation | Increased likelihood of predation at coastal haulouts; potential new marine predators | Low | Yes ($\phi$*_THM_*_)_ | N/A | N/A | Partially, through increased haulout mortality, but not modeled mechanistically | Data on haulout predation needed before mechanistic modeling can be done. Data needed on orca abundance and walrus predation in Bering and Chukchi Seas. |
| ANTHROPOGENIC DISTURBANCE STRESSORS |  |  |  |  |  |  |  |
| Oil & Gas Development^2^ | Loss in time spent foraging due to acoustic disturbance | Low-Moderately Low | Yes (*P_WATER_*, *P_FORAGE_*) | N/A | Yes | N/A | Intensity of oil & gas scenarios can be updated as more information on future energy development becomes available. |
| Commercial Fisheries | Displacement; increased energy expenditure; effect on prey abundance; by-catch | Moderately Low | Partially (*R*, *PD*) | N/A | N/A | Partially, through prey density, but not mechanistically | Loss in foraging time due to commercial fishing disturbance; studies on the relationship between commercial fishing and the density of walrus forage species |
| Shipping & Air Traffic^2^ | Displacement; Loss in time spent foraging due to acoustic disturbance; disturbance from coastal haulouts; direct strikes | Moderately Low-Moderate | Partially ($\phi$*_THM_*_)_ | N/A | N/A | Partially, through terrestrial haulout mortality. | Loss in foraging time due to acoustic disturbance from shipping and air traffic; estimates of future shipping traffic in Chukchi & Bering Seas. |
| Pollution | Increased likelihood of pollution from increased shipping and potential oil and gas development | Low-Moderately Low | No | N/A | N/A | N/A | Mechanistic response of walruses to an oil spill. |

^1^ Intensity classification given to winter/spring sea ice decline in the 2017 Species Status Assessment (MacCracken et al., 2017) was “Low-Moderately Low”; however, updated climate projections used in the present study suggest winter/spring sea ice loss will worsen.

^2^ These stressors were not included in the original MacCracken et al., (2017) table, but were added for this study and their intensity estimated based on summary statistics from their Bayesian Belief Network.

Table S2. Pacific walrus population parameters and vital rates used to calibrate the default DEB model.

| Parameter | Value (mean & 95% CrI) | Estimated Value in default DEB (mean & 95% CI) | Source |
| --- | --- | --- | --- |
| Adult Female Population Size (N) | 69,281 (43,242 – 100,259) | 69,281 (43,242 – 100,259) | Beatty et al., (2022) |
| N/K (Population Size / Carrying Capacity | 0.900 (1.000 with harvest)* | 0.888 (0.862 – 0.914) | MacCracken et al., (2017); Taylor et al., (2018) |
| Neonatal calf Survival  (age 0 – 3 months) | 0.900 (0.802 – 0.950) | 0.894 (0.891 – 0.897) | Taylor et al., (2018) |
| Older calf Survival  (age 3 months – 2 years) | 0.814 (0.699 – 0.923) | 0.784 (0.780 – 0.789) | Taylor et al., (2018) |
| Reproductive rate (annual probability a reproductive adult female gives birth to a female calf) | 0.125 (0.100 – 0.150) | 0.101 (0.100 – 0.102) | Taylor et al., (2018) |

*Note: N/K in 2015 is presumed to equal 1 (Taylor et al., 2018), with current harvest levels, and 0.9 without (MacCracken et al. 2017). We used the Pacific walrus theta-logistic model (Johnson et al., 2022) to estimate N/K in the absence of harvest.

Table S3. Estimates of daily haulout mortality rates for Pacific Walruses occupying terrestrial haulouts. These probabilities are used in a Bernoulli mortality trial for each day a walrus occupies a terrestrial haulout in the model, and they vary based on the region the walrus occupies.

| **Parameter** | **Region** | **Calves** | **Adults** |
| --- | --- | --- | --- |
| **Baseline Terrestrial Haulout Mortality** | | | |
| $\phi$*_THM_* | Alaska Chukchi (Regions 3 & 4) | 1.188 e^-04^ | 4.808 e^-05^ |
| $\phi$*_THM_* | Russia Chukchi (Regions 1 & 2) | 3.175 e^-04^ | 8.788 e^-05^ |
| $\phi$*_THM_* | Bering Sea (Region 0) | 5.817 e^-04^ | 8.270 e^-06^ |
| **“Bad Year” Terrestrial Haulout Mortality** | | | |
| $\phi$*_THM_BY_* | Alaska Chukchi (Regions 3 & 4) | 7.503 e^-04^ | 2.006 e^-04^ |
| $\phi$*_THM_BY_* | Russia Chukchi (Regions 1 & 2) | 1.395 e^-03^ | 8.788 e^-05^ |
| $\phi$*_THM_BY_* | Bering Sea (Region 0) | 1.055 e^-04^ | 1.499 e^-05^ |

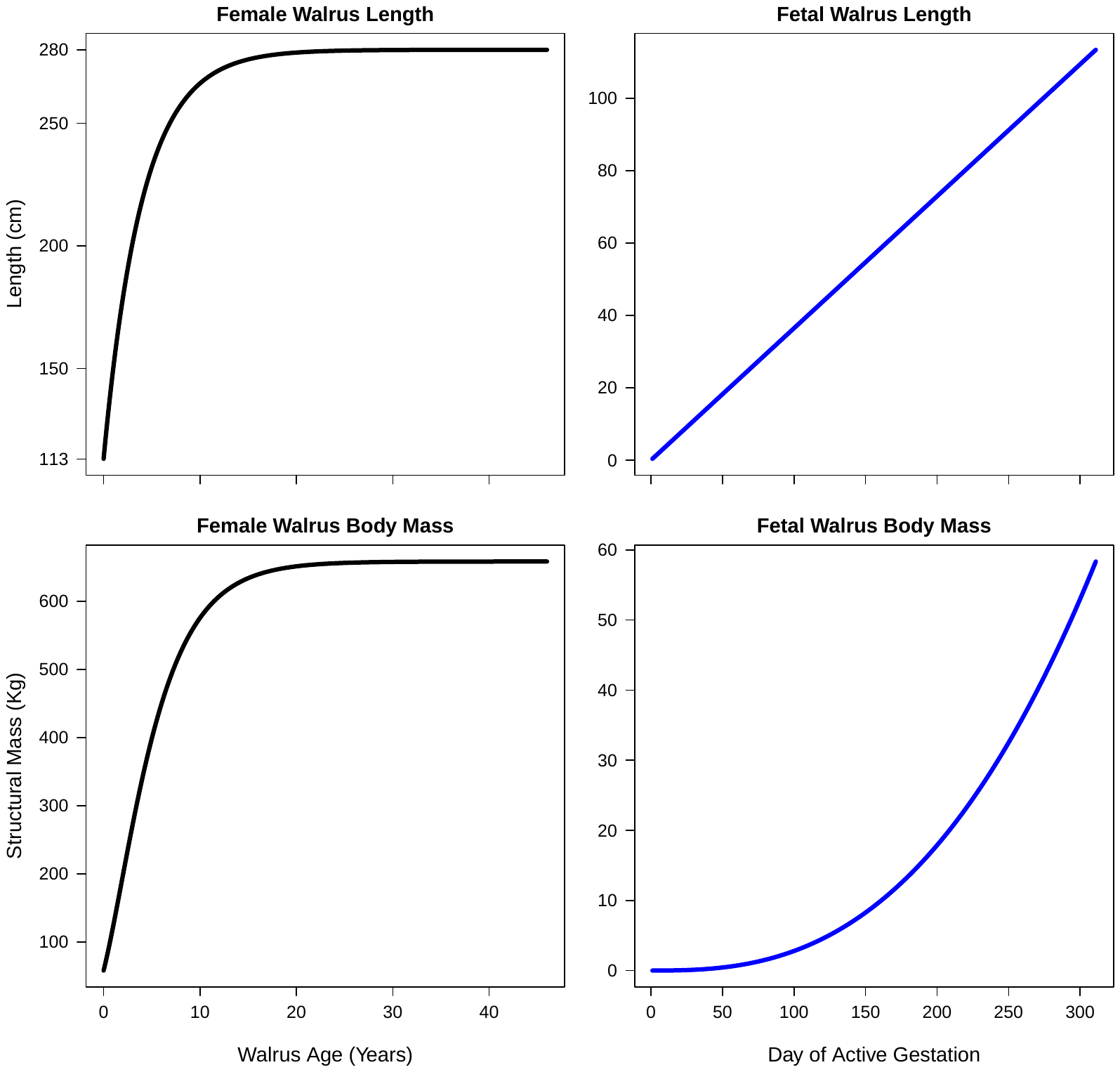

Figure S1. Estimated growth curves (length and structural body mass (i.e., non-reserve mass)) for female Pacific walruses and their fetuses. Growth curves were used to estimate the energetic costs of growth and fetal maintenance and determine a female’s body size at a given age.

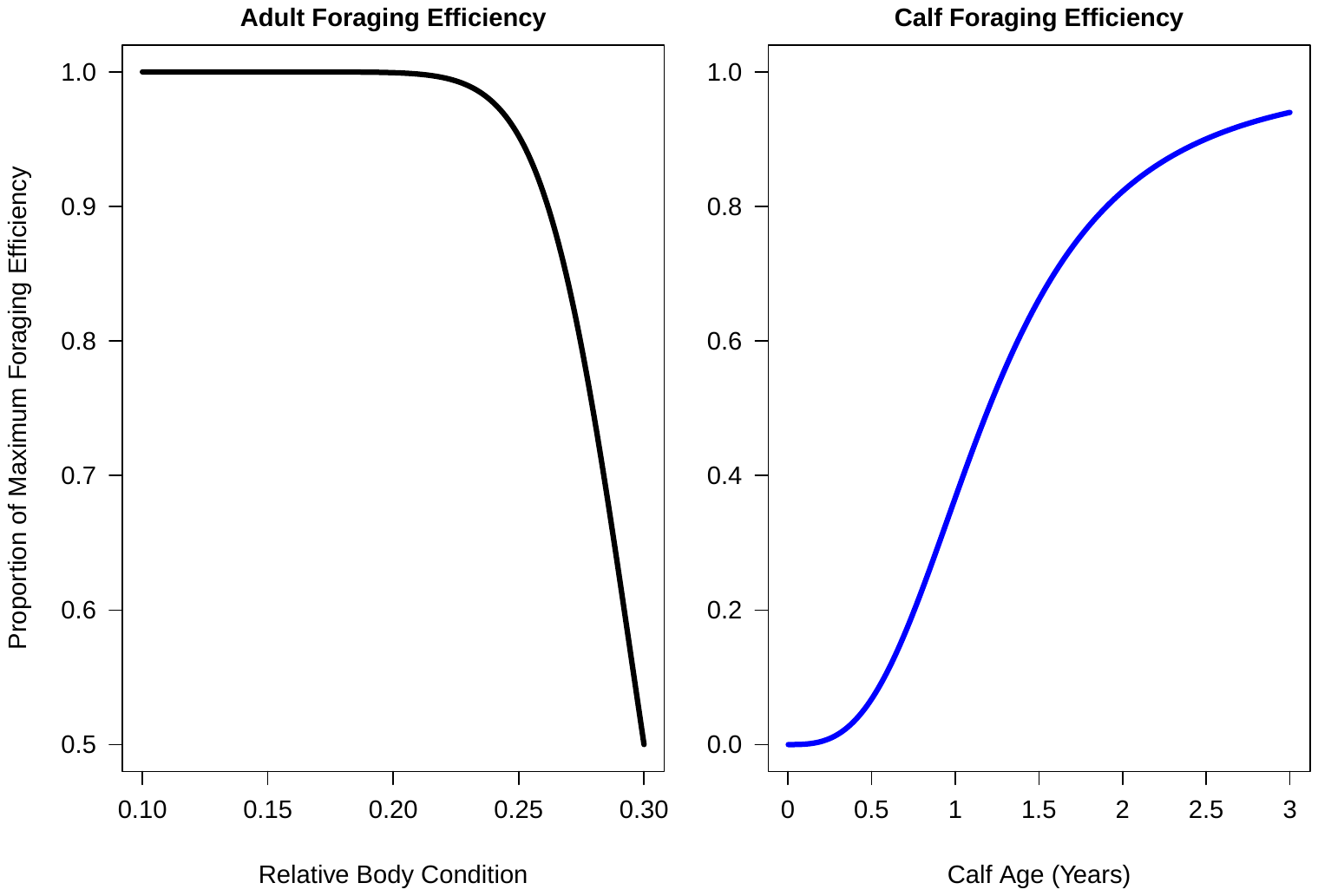

Figure S2. The assumed relationship between foraging efficiency and body condition (i.e., reserve mass/total mass) for adult Pacific walruses and between foraging efficiency and age for Pacific walrus calves from 0-3 years of age used in the DEB.

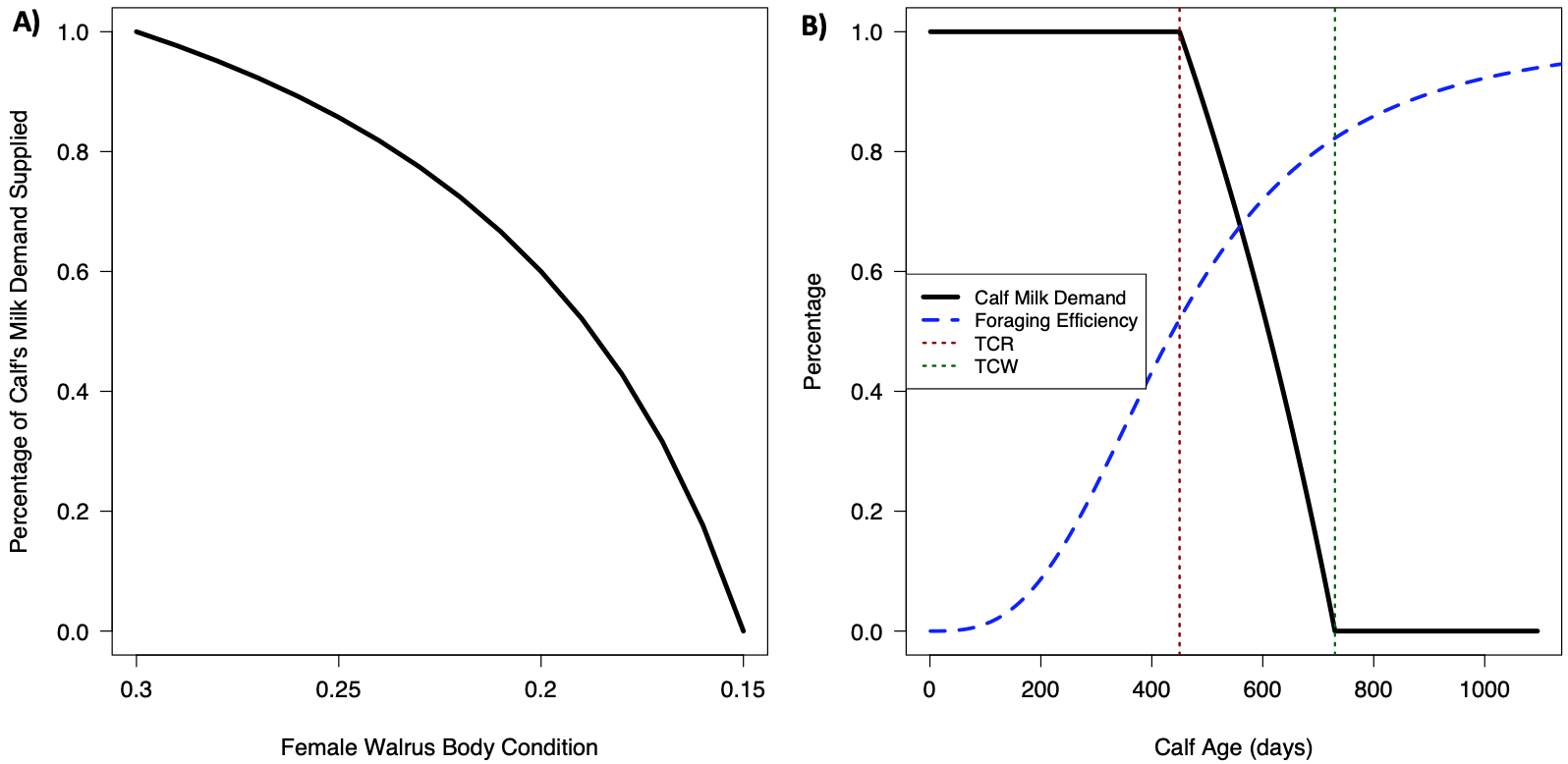

Figure S3. The assumed relationship between female Pacific walrus body condition (i.e., reserve mass/total mass) and the percentage of her calf’s milk demand she is able to supply (*ψ_t_*, panel A), and the relationship between calf age and its milk demand (the percentage of its energy demand that comes from milk; Panel B). TC_R_ (450 days) is the age at which the female begins to reduce milk supply to the calf, and TC_W_ (730 days) is the calf’s age at weaning.

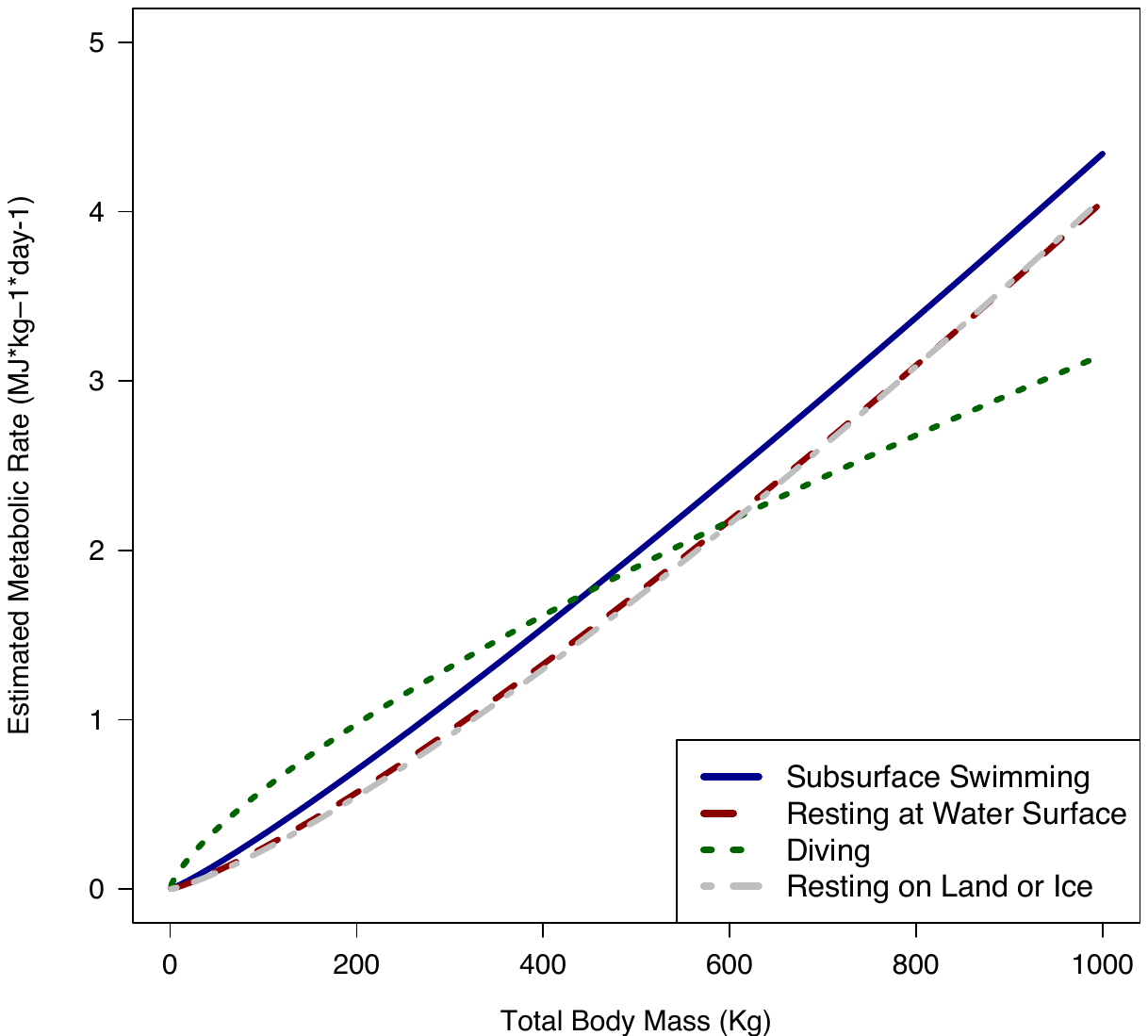

Figure S4. Mass-specific metabolic rates for the four walrus activity states considered in the Dynamic Energy Budget. Details on the associated equations can be found in Table 1 of the main article.

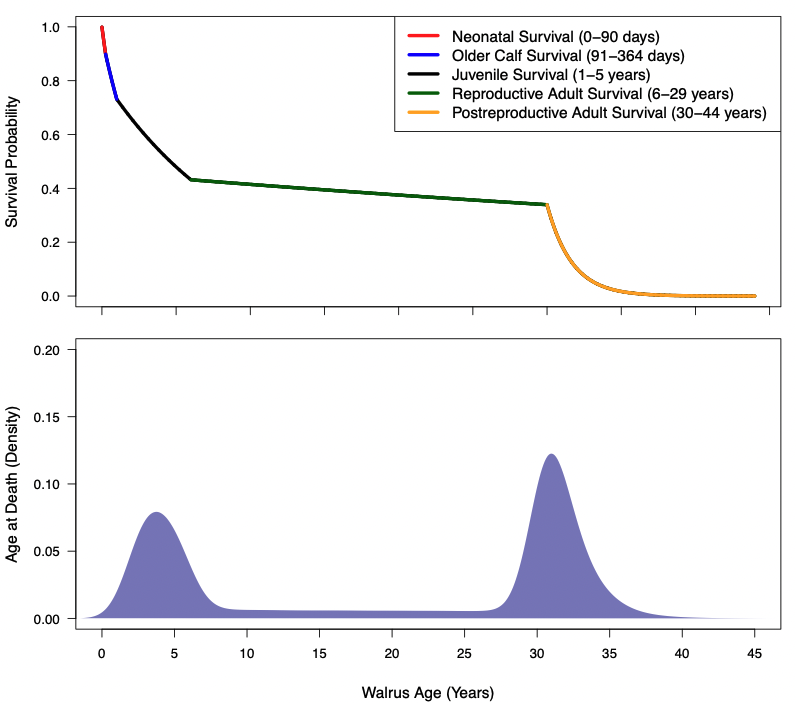

Figure S5. Cumulative survival curve for the Pacific walrus (top panel), and probability density of age at death (bottom panel). Age-specific annual survival rates were based on Taylor et al. (2018). This results in a bimodal distribution of simulated age at death, with one peak during elevated juvenile mortality and another during elevated post-reproductive mortality.

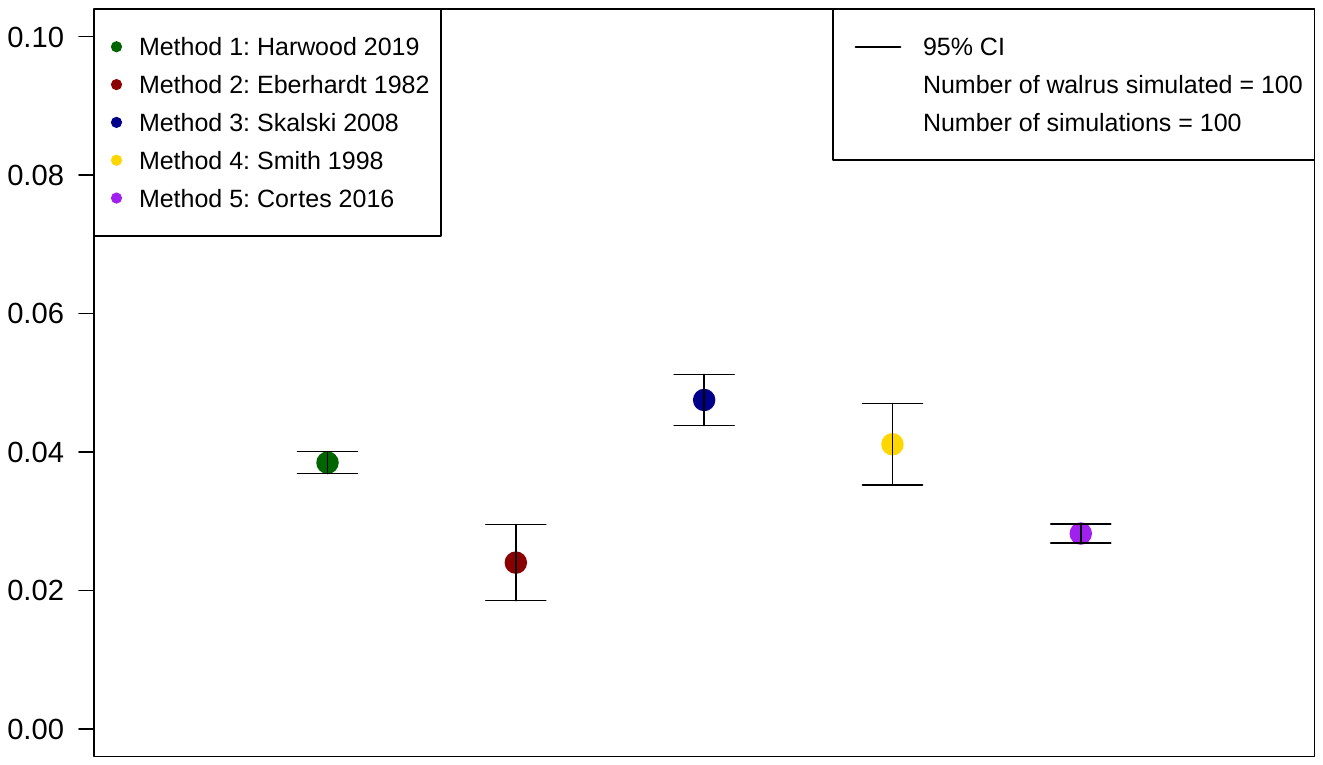

Figure S6. Intrinsic rate of increase (*r_max_*) estimates from the method applied in the current study (Method 1) and five additional methods discussed in Cortes (2016). All information used to calculate *r_max_* (e.g., survival and annual reproductive rate) were estimated using the DEB calibrated to baseline parameter values (i.e., using the values in Table S2).

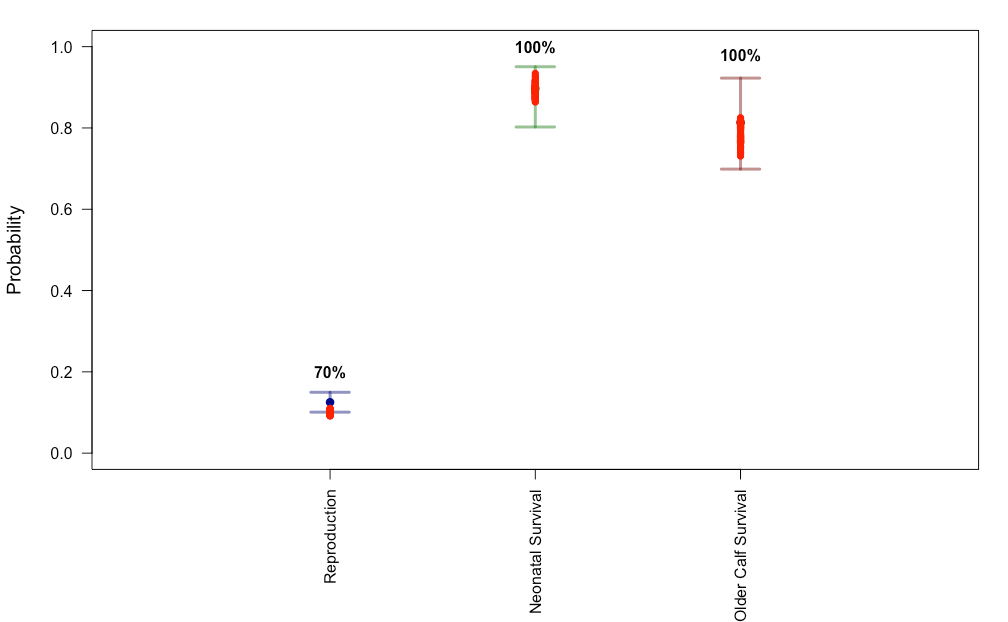

Figure S7. Calibration of the DEB to density dependent rates (annual reproductive rate [# female calves/reproductively mature female per year], neonatal and older calf survival) from the Taylor et al. (2018) integrated demographic population model (IPM) most parsimonious model, using an *R* of 4.034 and a *φ_M_* of 4.034. Colored bars indicate the 95% credible interval of vital rates in the year 2015 from the IPM, and red points are 100 DEB simulations of 100 individuals. Percentages indicate the percent of simulations that fell within the 95% CrI for each parameter.

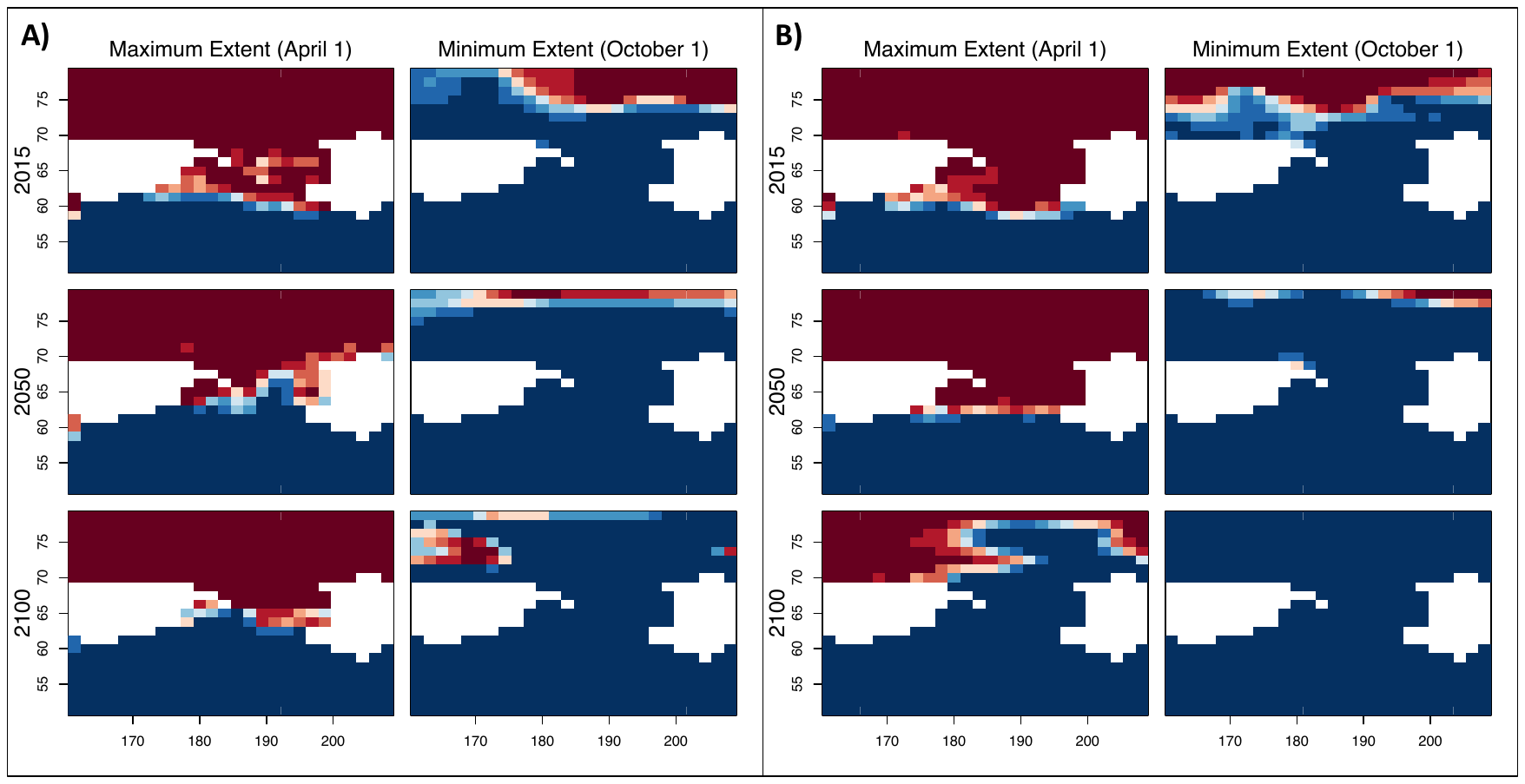

Figure S8. Example of CMIP6 sea ice model output [ACCESS-ESM1-5] for A) ssp245 (intermediate) and B) ssp585 (pessimistic) models. Panes portray the maximum and minimum sea ice extent (standardized to April 1 and October 1, respectively) for 2015, 2050, and 2100 in the Bering and Chukchi seas. Blue is water, maroon is ice, and white is land, with Russia on the left and Alaska on the right. The x-axis gives longitude, the y-axis gives latitude, and the Bering and Chukchi seas are delineated at a latitude of 66°N.

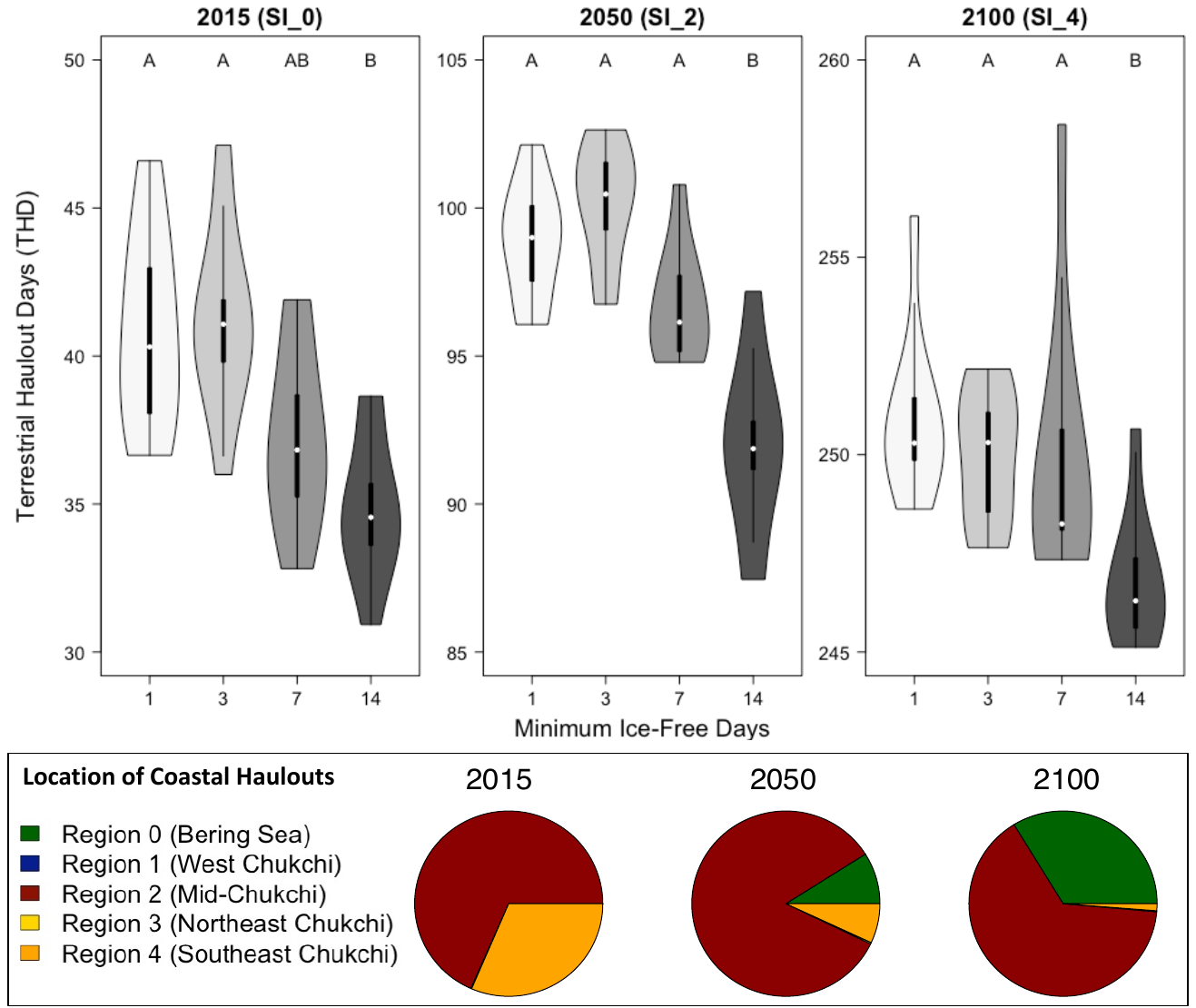

Figure S9. Estimates of the number of terrestrial haulout days per year (THD) for 2015 (SI_0), 2050 (SI_2) and 2100 (SI_4 sea ice scenario). Upper graph shows sensitivity of the THD (aggregated across regions) to the number of days of ice-free conditions (1, 3, 7, or 14) before a simulated walrus must rest at a terrestrial haulout. Simulations were conducted on 100 populations of 100 individuals for each sea ice scenario and ice-free day group. Letters represent significance groups from an ANOVA and post-hoc Tukey test. Pie charts indicate the regions where simulations estimate the terrestrial haulout days occur following the movement patterns outlined in Udevitz et al. (2017). Note different scales on the y-axes.

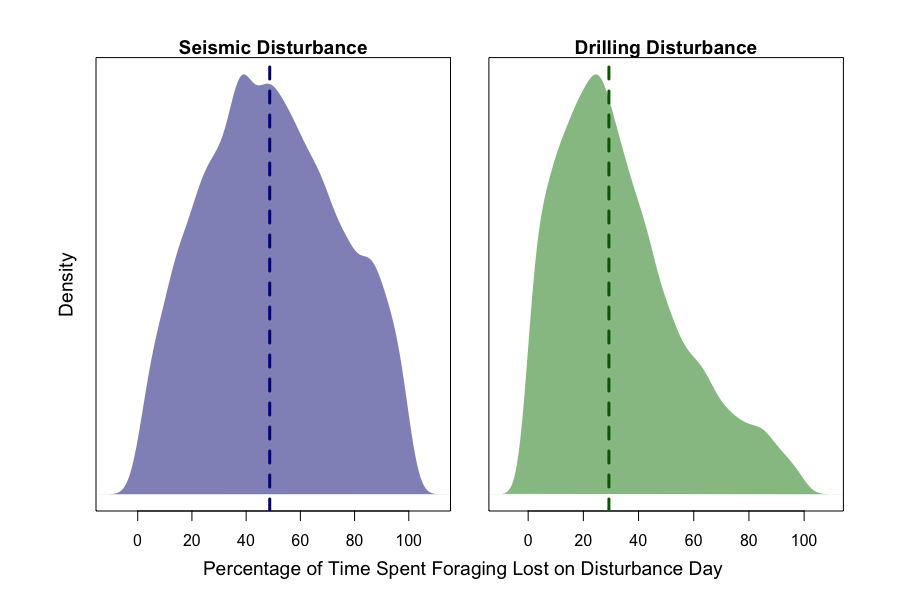

Figure S10: Results of an Expert Elicitation (EE) linking the behavioral response of Pacific walruses to disturbance associated with oil and gas activities (from Harwood et al., 2019). Vertical lines indicate the medians of the associated probability distributions.

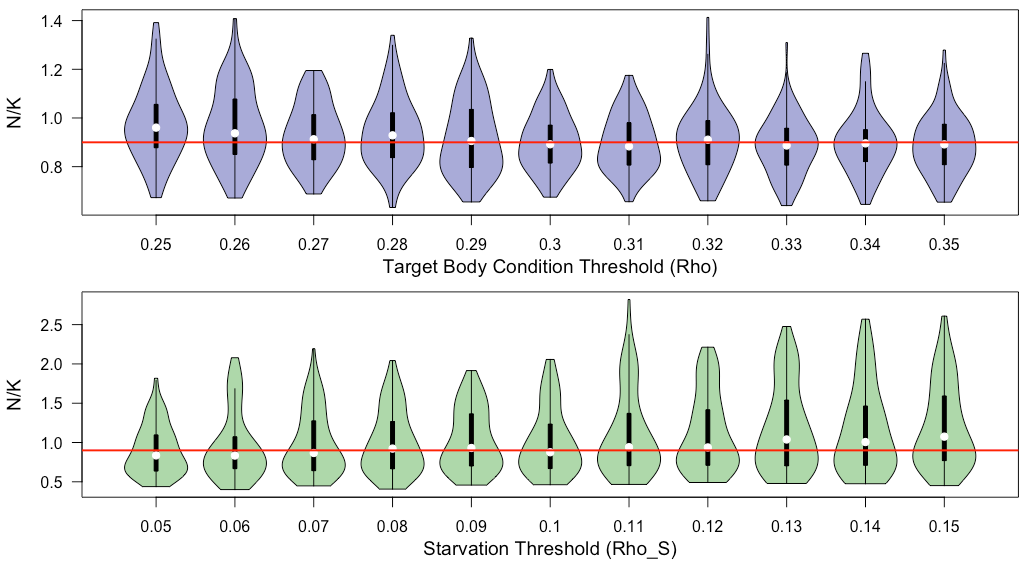

Figure S11. Sensitivity of *N/K* (population size/carrying capacity) to different values of *ρ* (the target body condition) and *ρ_s* (the starvation threshold). Red horizontal lines indicate an *N/K* value of 0.9, which was the target value for model calibration. We conducted 100 simulations, each comprised of 100 individuals for each body condition value, and there were no significant differences for the values considered based on an ANOVA and post-hoc Tukey test. Note different scales on y-axes.

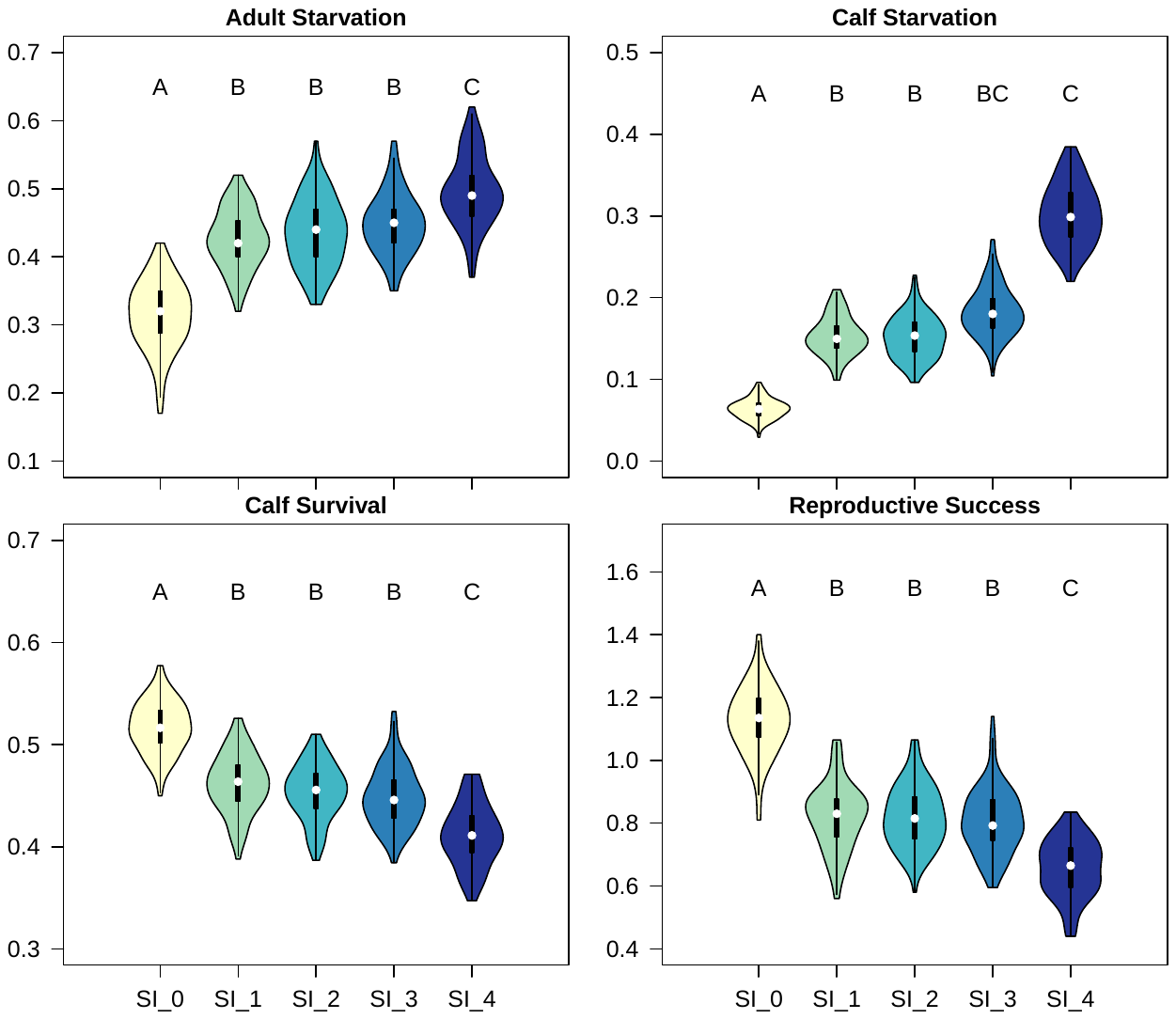

Figure S12. Sensitivity of four demographic parameters to the five sea ice scenarios (SI_0–SI_4, which represent progressively lessening ice cover) modelled in the DEB, considered independently from terrestrial haulout mortality. For each sea ice scenario, we conducted 100 simulations each of 100 individuals. Letters represent significance groups from an ANOVA and post-hoc Tukey test. Adult starvation is the probability of an adult dying of starvation over the course of its life; calf starvation is the number of calves that die of starvation per simulated female; calf survival is the probability of a calf surviving the first two years of its life; and reproductive success is the average number of female calves that survive to weaning per each mature female for each simulation. Note different scales on y-axes.

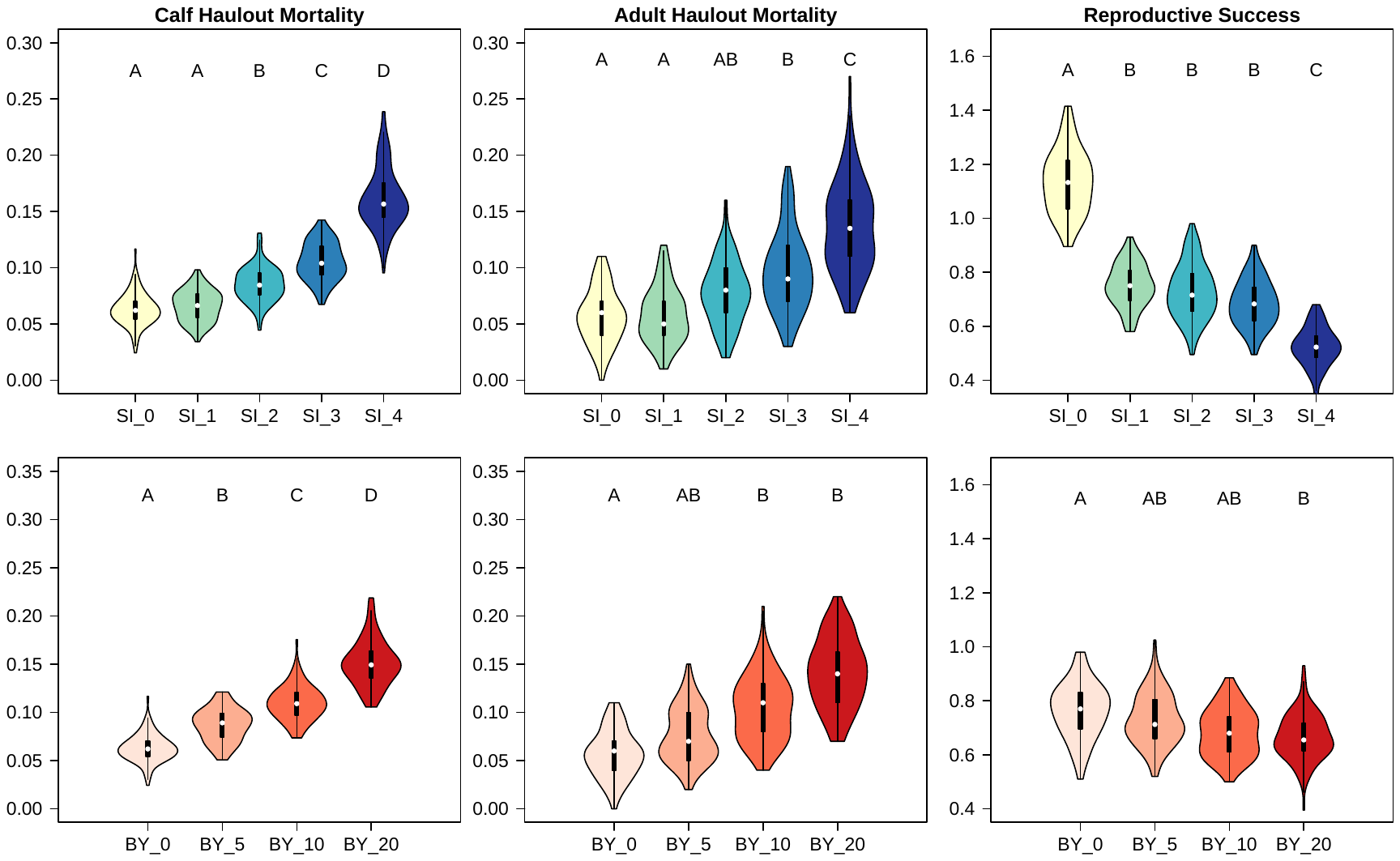

Figure S13. Sensitivity of three DEB demographic parameters to terrestrial haulout mortality under the five sea ice scenarios considered in this study (top panels; SI_0–SI_4), and the effects of changing the “bad year” (BY) haulout parameter in the model to 0, 5, 10, and 20 years per each simulated walrus’ lifetime (bottom panels). Top panels show the effect of haulout mortality with a BY value of 0. Simulations in bottom panels hold all other parameters constant under a SI_1 sea ice scenario to demonstrate the effect of various BYs in isolation. Calf haulout mortality is a calf’s probability of dying at a haulout in the first two years of its life; adult haulout mortality is the simulated female’s probability of dying at a haulout over the course of its life; and reproductive success is the average number of female calves that survive to weaning per each mature female for each simulation. These models include other sources of mortality (e.g., starvation) associated with sea ice availability. For each sea ice scenario and BY group, we conducted on 100 simulations each of 100 individuals. Letters represent significance groups from an ANOVA and post-hoc Tukey test. Note different scales on y-axes.

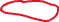

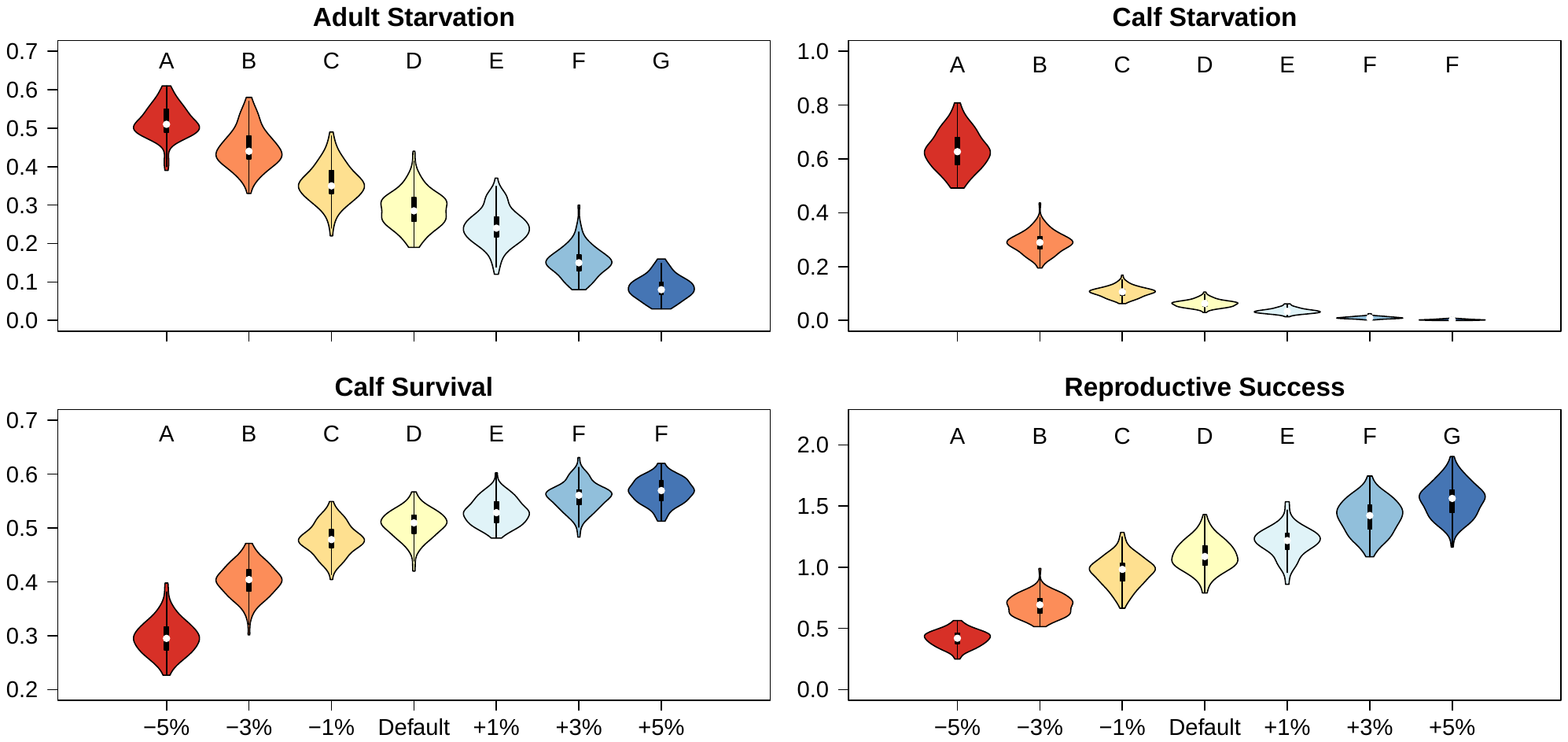

Figure S14. Sensitivity of DEB outcomes to adjusting resource availability (*R*) by ± 1–5% from the baseline scenario. We conducted 100 simulations each of 100 individuals, and letters represent significance groups from an ANOVA and post-hoc Tukey test. Adult starvation is the probability of an adult dying of starvation over the course of its life; calf starvation is the number of calves that die of starvation per simulated female; calf survival is the probability of a calf surviving the first two years of its life; and reproductive success is the average number of female calves that survive to weaning per each mature female for each simulation. Note different scales on y-axes.

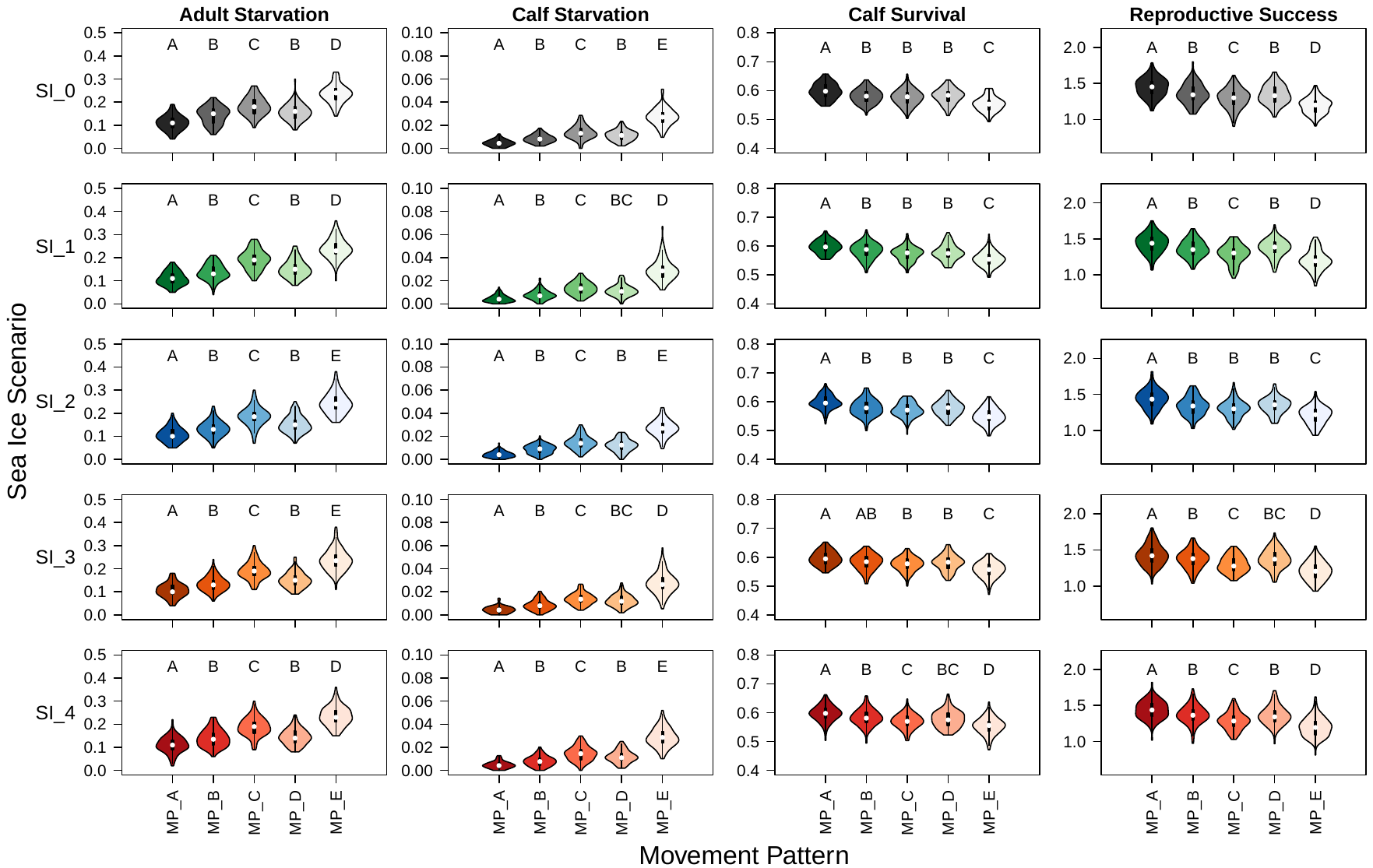

Figure S15. Sensitivity of DEB outcomes to movement pattern. Each row portrays a different sea ice scenario (i.e., SI_0-SI_4), each column displays a different outcome, and each plot is one of the five primary movement patterns (A-E) in isolation. We conducted 100 simulations each of 100 individuals. Letters represent significance groups from an ANOVA and post-hoc Tukey test. Adult starvation is the probability of an adult dying of starvation over the course of its life; calf starvation is the number of calves that die of starvation per simulated female; calf survival is the probability of a calf surviving the first two years of its life; and reproductive success is the average number of female calves that survive to weaning per each mature female for each simulation. Note different scales on y-axes. In general, movement patterns confined to the eastern Chukchi Sea conferred higher reproductive success and calf survival, and lower rates of starvation. The movement pattern that conferred the lowest rates of reproductive success and calf survival and the highest starvation rates not only extended across the eastern and western Chukchi Sea, but it also did not extend nearly as far north as any of the other patterns.

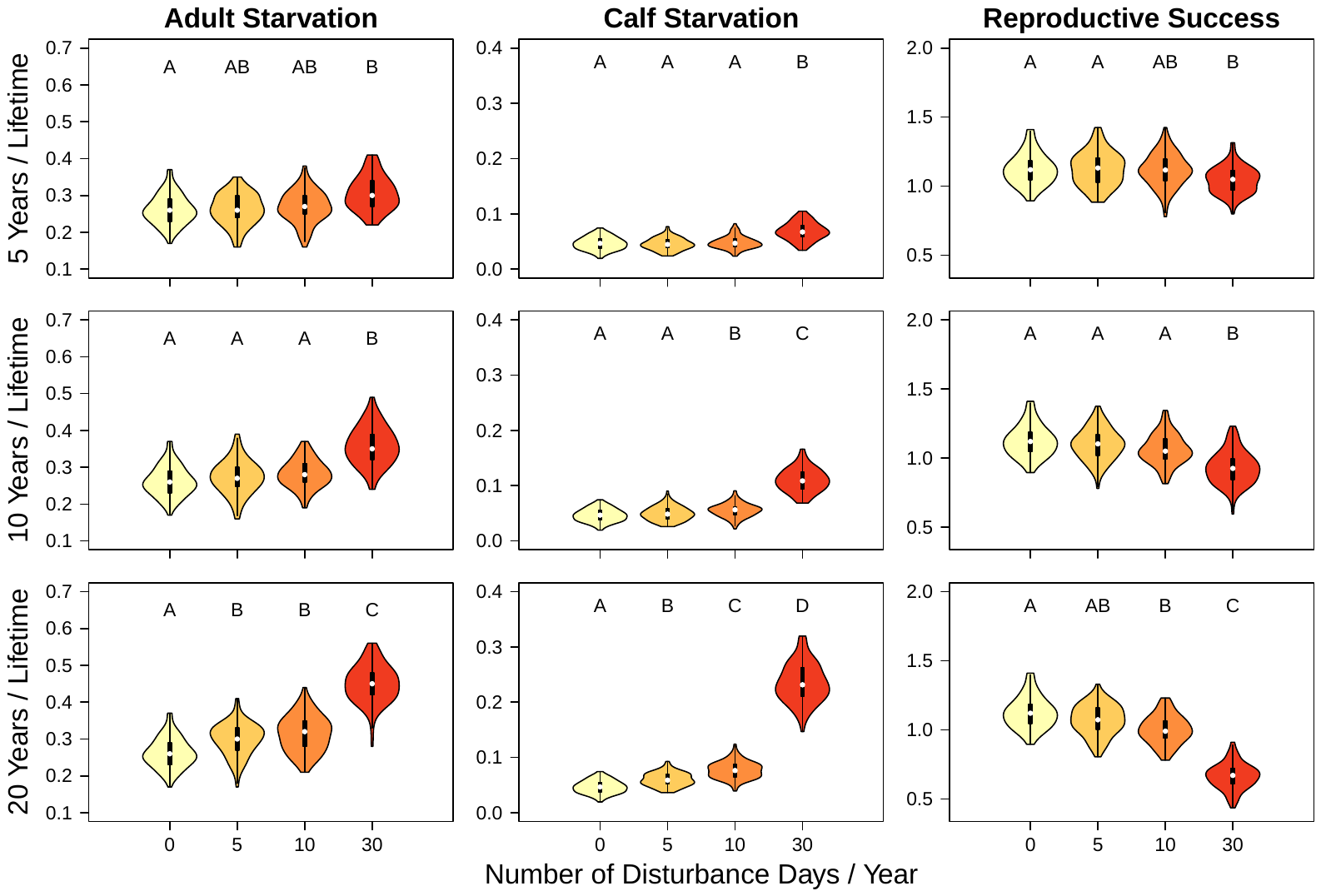

Figure S16. Sensitivity of DEB outcomes to anthropogenic disturbance. Simulated walruses in the baseline model were subjected to 0, 5, 10, and 30 randomized disturbance days/year (columns, colors) for 5, 10, and 20 randomized years/lifetime (rows). Each disturbance day was randomly assigned either the “seismic” or “drilling” disturbance type. We conducted 100 simulations each of 100 individuals. Letters represent significance groups from an ANOVA and post-hoc Tukey test. Adult starvation is the probability of an adult dying of starvation over the course of its life; calf starvation is the number of calves that die of starvation per simulated female; calf survival is the probability of a calf surviving the first two years of its life; and reproductive success is the average number of female calves that survive to weaning per each mature female for each simulation. Note different scales on y-axes. In general, human disturbance had the strongest effect when each walrus was exposed to 30 days of disturbance/year, and this effect increased as walruses had more disturbed years throughout their simulated lifetimes. In general, increasing disturbance to 30 days/year had the most significant effect on simulated walruses, and increasing the number of years/lifetime that disturbance occurs compounded that effect.
